# *SNCA*-targeted epigenome therapy for Parkinson’s disease alleviates pathological and behavioral perturbations in a mouse model

**DOI:** 10.1101/2025.06.16.659948

**Authors:** Bernadette O’Donovan, Joseph Rittiner, Suraj Upadhya, Dellila Hodgson, Boris Kantor, Ornit Chiba-Falek

**Author notes:** Corresponding authors: Ornit Chiba-Falek, Division of Translational Brain Sciences, Department of Neurology, Duke University School of Medicine, Durham, North Carolina 27710, USA, Boris Kantor, Department of Neurobiology, Duke University School of Medicine Durham, North Carolina 27707, USA.

## Abstract

Alpha-synuclein (SNCA) overexpression is implicated in Parkinson’s disease (PD) pathogenesis, making SNCA downregulation a promising therapeutic strategy. We developed a *SNCA*-targeted epigenome therapy using an *all-in-one* lentiviral vector (LV) carrying deactivated CRISPR/(d)Cas9, gRNA targeted at *SNCA*-intron1, and either the catalytic domain of DNA-methyltransferase3A (DNMT3A), or a synthetic repressor molecule of Krüppel-associated box (KRAB)/ methyl CpG binding protein 2 transcription repression domain (MeCp2-TRD). Therapeutic efficacy was evaluated in a new PD mouse model, generated with an adeno-associated viral vector carrying an engineered minigene comprised of the human (h)A53T-*SNCA* expressed via the human native regulatory region. Both therapeutic vectors reduced expression of α-synuclein in the substantia nigra (SN), with LV/dSaCas9-KRAB-MeCP2(TRD) demonstrating greater repression. LV/dSaCas9-KRAB-MeCP2(TRD) also significantly reduced pathological α-synuclein aggregation and phosphorylation (Ser129), and preserved tyrosine hydroxylase expression in the SN and the striatum. Behavioral analysis following LV/dSaCas9-KRAB-MeCP2(TRD) injection, showed significant improvement in motor deficits characteristic of our PD-mouse model. Safety assessments found normal blood counts, serum chemistry, and weights. Collectively, we provide *in vivo* proof-of-concept for our *SNCA*-targeted epigenome therapy in a PD-mouse model. Our results support the system’s therapeutic potential for PD and related synucleinopathies and establish the foundation for further preclinical studies toward investigational new drug enablement.

## INTRODUCTION

Parkinson’s disease (PD) is the second most common neurodegenerative disorder. Clinically, it is a movement disease, characterized by resting tremor, bradykinesia, rigidity, and postural instability ^1^. The neuropathological hallmarks of PD include neuronal loss primarily in the substantia nigra (SN) and abnormal protein aggregates, known as Lewy bodies (LBs) and Lewy neurites ^2–6^ composed largely of the insoluble α-synuclein protein (α-syn) ^7^. *SNCA* has been well established as a major genetic factor of both the familial and sporadic forms of PD. Several autosomal-dominant mutations in the *SNCA* gene have been identified in families with the rare early onset PD ^8–17^. Genome wide association studies (GWAS) have repeatedly identified *SNCA* as a significant risk factor for the common non-Mendelian form of PD ^18–33^. Consequently, *SNCA* is one of the most promising therapeutic targets for PD.

Accumulating studies have established that elevated levels of wild type α-syn are causative in the pathogenesis of synucleinopathies (reviewed in ^34^). Firm evidence was provided by the identification of *SNCA* multiplications in individuals with autosomal-dominant early-onset parkinsonism and rapid cognitive decline ^35–38^. These studies reported a dose-dependent effect of *SNCA* levels on disease manifestation, *i.e.,* the triplication caused an earlier age at onset and a more severe clinical phenotypes than the duplication ^36^. Moreover, higher *SNCA* expression levels were described in post-mortem brain tissues from sporadic PD patients compared to healthy controls using bulk ^39–43^ and single-cell analyses ^44^. In addition to human studies, cell-based and animal models supported the causative role of *SNCA* overexpression in the development of PD and related disorders (*34*). Consistently, previous studies in various rodent models of PD showed that lowering α-syn levels had neuroprotective effects ^45–47^. While these were promising results, others reported that robust α-syn knockdown mediated by RNAi led to nigrostriatal degeneration in both rodents and non-human primates ^48–51^. Collectively, these studies suggested that the reduction in α-syn levels has a therapeutic window within which beneficial effects will be achieved without disruption of normal neuronal function. Thus, fine-tuning *SNCA* gene expression vs. robust knockdown presents a strategy for the development of disease modifying therapy (DMT) for PD and related synucleinopathies.

*SNCA* gene expression is regulated by various mechanisms including epigenetic modification ^52^. Demethylation of a CpGs island within *SNCA* intron 1 in post-mortem brain tissue and blood has been associated with PD and was correlated with increased *SNCA* expression ^53–58^. This data defined *cis*-regulatory elements within *SNCA* intron 1 that impact the regulation of *SNCA* expression in the context of PD. We have translated this knowledge into the development of tools to reduce *SNCA* expression levels accurately and specifically via epigenome editing targeted at those regulatory sequences within *SNCA* intron. Previously we demonstrated the specific repression of α-syn levels in human induced pluripotent stem cell (hiPSC) derived dopaminergic neurons from a PD/dementia with Lewy body disease (DLB) patient with the *SNCA* triplication, that was sufficient to ameliorate disease associated perturbations ^59, 60^. Here, we advanced our *SNCA*-targeted epigenome-editing system into *in vivo* validation in a mouse model of PD. Towards this goal, we utilized a novel PD mouse model based on adeno-associated virus (AAV)-induced overexpression of the human (h)A53T-α-syn. A53T SNCA is a variant of SNCA associated with familial forms of PD that is thought to promote misfolding and aggregation of α-syn ^16, 61, 62^. The vector comprised of a minigene carrying the coding sequence of the A53T mutated human-*SNCA* fused with the human native promoter/intron 1 region.

Furthermore, we compared *in vivo*, in the mouse SN, the efficacies and efficiencies of the two transcription repressor molecules, *i.e.,* the catalytic domain of DNA-methyltransferase 3A (DNMT3A) and the synthetic Krüppel-associated box (KRAB)/methyl CpG binding protein 2 (MeCP2) transcription repression domain (TRD). Our study validated *in vivo* the target engagement, efficacy and specificity of our *SNCA*-targeted epigenome-editing system. We demonstrated the applicability of the system to reduce α-syn expression levels in the SN of our PD mouse model, and its utility in reversing disease associated pathological hallmarks and motor impairment characteristics of the PD model.

## RESULTS

### Vector design and in vitro proof-of-concept

We engineered a reporter vector expressing *SNCA* open reading frame (ORF) fused with destabilized GFP (dGFP) and nano-luciferase (NLuc) genes that are characterized by short protein half-lives, suitable for the evaluation of gene expression changes. We used CAG promoter harboring *SNCA* intron 1 sequence to express this vector and a lentiviral backbone harboring puromycin selection marker ^59^ (Fig. 1a). The inactive mutants of Cas9 protein, derived from *Staphylococcus aureus* (dSaCas9) (Fig. 1a) has been linked with a Krüppel associated box (KRAB) domain from Zinc finger protein 10 (KOX1), and a transcription repressor domain from MeCP2 (TRD) to create a KRAB-MeCP2 bimodal repressor ^63^, or fused to the catalytic domain of DNA methyltransferases 3A (DNMT3A) ^59^ (Fig. 1a). For each repressor molecule a unique gRNA was selected to target the reporter construct in the h*SNCA* promoter/intron1 region (Fig. 1a), and a luciferase assay was performed. In HEK293T reporter cells we did not observed reduction in NLuc expression for samples transduced with vectors not expressing a CAG/intron1-targeting gRNA (across all effectors, including a no-gRNA/no-effector double control; Fig. 1b). Evaluation of the effects of the selected DNMT3A and KRAB-MeCP2 molecules demonstrated significant levels of repression (Fig. 1b). Nevertheless, LV/dSaCas9-KRAB-MeCP2(TRD), demonstrated a substantial greater silencing, *i.e.,* a reduction of NLuc expression by about 85% vs ∼ 40% (Fig. 1b). The KRAB-MeCP2(TRD) vector with gRNA2 and DNMT3A vector with gRNA1 showed no noticeable repression of the reporter construct (Fig. 1b). Noteworthy, gRNA1 targets a sequence which is proximal to the transcription start site (TSS), whereas gRNA2 targets an upstream part of the promoter (Fig. 1a), thus, these results are consistent with earlier findings demonstrating that higher levels of repression are often achieved when gRNAs are designed to target a sequence in the vicinity of the TSS ^64, 65^. Importantly, the results of DNMT3A-gRNA2-mediated repression are consistent with our earlier observations ^59^.

**Figure. 1.**
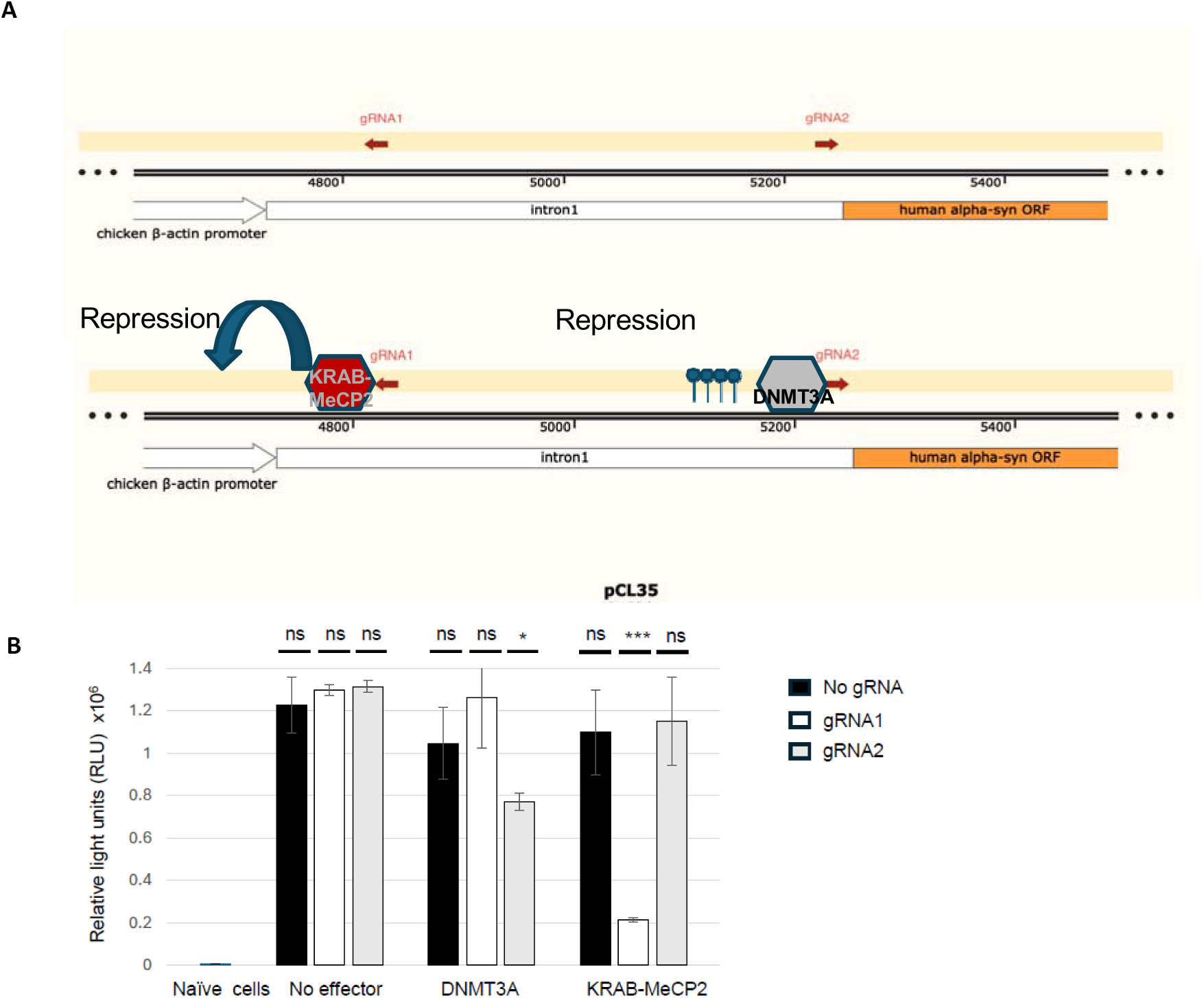
In vitro validation of dSaCas9-DNMT3A and dSaCas9-KRAB-MeCP2(TRD) (**A**) Schematic showing reporter vector expressing alpha-synuclein ORF fused with destabilized GFP (dGFP) and nano-luciferase (NLuc) genes. To express this cassette, we used CAG promoter harboring SNCA intron 1 sequence. dSaCas9 has been linked with a KRAB-MeCP2(TRD) bimodal repressor, or fused to the catalytic domain of DNMT3A. Two gRNAs sequences were selected to target the reporter construct in the promoter/intron1 region. (**B**) The vectors were transduced into the HEK293T reporter cell line to evaluate their repressive effects. The cells were harvested at day 4 post transduction, total protein was normalized via BCA assay, and a luciferase assay was performed. No reduction in NLuc expression was detected in any of the samples transduced with vectors not expressing a CAG/intron1-targeting gRNA. Both DNMT3A and KRAB-MeCP2(TRD) constructs showed significant levels of repression. dSaCas9-KRAB-MeCP2(TRD), demonstrated the most substantial silencing by a large margin. * p < 0.05 *** p < 0.001.

### AAV-induced hA53T *SNCA* mouse model for PD

For *in vivo* proof-of-concept studies we developed a humanized PD mouse model induced by overexpression of the hA53T-α-syn expressed from an engineered minigene. The minigene of this model was developed based on the published AAV vector ^66^, including the human noncoding sequences within the promoter/intron 1 of region of h*SNCA* to imitate the expression pattern in the human brain. The vector comprised of a minigene carrying the coding sequence of the A53T mutated human-*SNCA* fused with the human native promoter/intron 1 region (Fig. 2a). We characterized the model using twelve C57BL/6 mice (6 male, 6 female) stereotaxically injected with the AAV/hA53T-*SNCA* vector (1 µL; 1×10^13^ vg/mL) into the SN in one hemisphere and compared to a null AAV vector injected into the other SN (1 µL), such that each mouse provided an internal control (Fig. 2b). There was no significant difference in expression between males and females, thus, the data was combined for subsequent analysis. Immunohistochemistry analysis 6 weeks post injection revealed that the AAV/hA53T-*SNCA* vector induced overexpression of total human α-syn (46-fold higher than null AAV, t(11)=4.9 p=0.0005, paired t-test), and PD like pathology showing higher levels of pSer129 α-syn (29-fold higher than null AAV, t(11)=2.98 p=0.0125, paired t-test), and human α-syn positive aggregates (Fig. 2c-d). To evaluate the impact on dopaminergic neurons, we assessed endogenous mouse tyrosine hydroxylase (TH) expression in the SN. Observation of a 38% decrease in TH expression indicated a loss of dopaminergic neurons in the SN (t(11)=18.75 p=0.0001, paired t-test, Fig. 2c,e). Injection of the AAV/hA53T-*SNCA* vector also induces motor deficits as measured by a cylinder test to assess forepaw function. Following unilateral injection of AAV/hA53T-*SNCA* into the SN, significant asymmetry of forepaw use was observed with preference for the side ipsilateral to injection site (see Fig. 5). These data showed that injection of AAV/hA53T-*SNCA* into the mouse SN leads to human α-syn protein overexpression and aggregation, decreases in TH positive neurons, and motor deficit, thereby modeling pathological hallmarks of PD.

**Figure 2.**
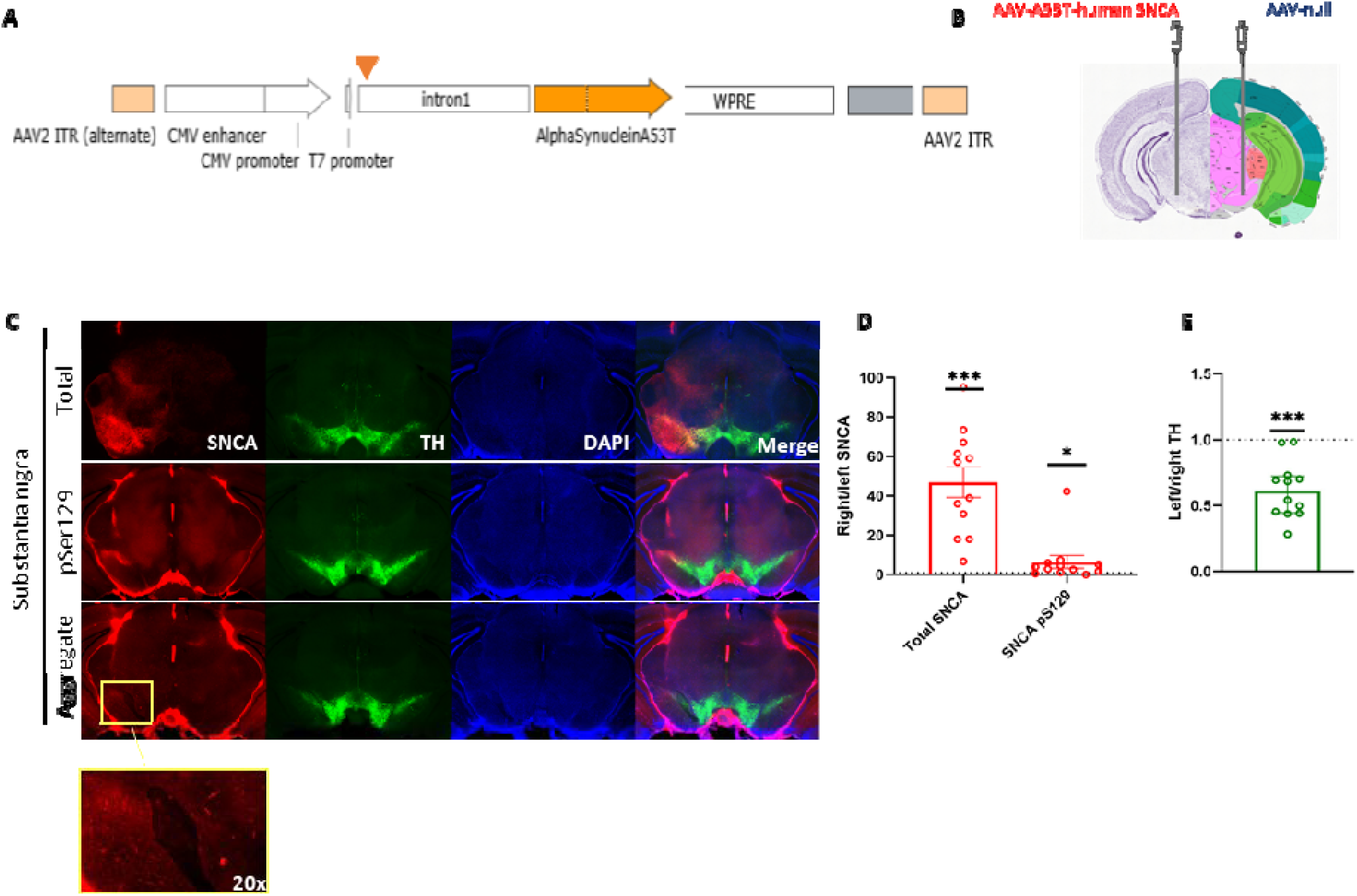
AAV/A53T-human SNCA induced mouse model for PD. (**A**) Schematic showing the structure of the AAV/A53T-human SNCA vector. (**B**) Diagram of experimental design; the AAV/A53T-human SNCA vector was stereotactically injected into the SN in one hemisphere and a null AAV vector into the other SN. Image modified from the Allen Brain Atlas (allenbrainatlas.org). (**C**) Representative images of brain coronal slices at 2x magnification 6 weeks post-injection, showing protein expression of human total α-syn, α-syn pS129, and aggregated α-syn, and mouse TH, and DAPI staining in the SN. α-syn aggregates were observed in the SN induced by the AAV-A53T-human SNCA vector (yellow box). Fluorescence intensity was quantified using ImageJ. Box plot displays the ratios of the right SN relative to the left SN. Each open circle represents the quantified signal right/left for a mouse. (**D)** AAV-A53T-human SNCA vector induced strong expression of human total α-syn and α-syn pS129 in the SN. (**E**) Expression of TH was significantly lower in the right SN relative to the left SN. Values represent mean ± SEM. N =12 (6 female, 6 male) * p < 0.05 *** p < 0.001; Paired t-test.

### LV/dSaCas9-DNMT3A reduces α-synuclein expression in PD mouse model

Previously, we reported that a LV/dCas9-DNMT3A system successfully downregulates α-syn mRNA and protein, and rescued disease related cellular phenotypes in hiPSC-derived dopaminergic neurons from a PD/DLB patient with the *SNCA* triplication ^59^. To investigate the *in vivo* efficacy of the LV/d*Sa*Cas9-DNMT3A therapeutic system (Fig. 3a), a total of ten C57B/L6 mice (4 males, 6 females) received bilateral stereotaxic injections. AAV/hA53T-*SNCA* (2.5×10^9^ vg) was co-injected with LV/dSaCas9-DNMT3A (7.5×10^6^ vg) into the right SN of mice, and the same amount of AAV/hA53T-*SNCA* was co-injected with the LV/dSaCas9 control (7.5×10^6^ vg) (no repressor, no gRNA) into the left SN (Fig. 3b). At six weeks post-injection, the expression of total human α-syn, α-syn pS129, and aggregated α-syn was quantified by immunohistochemistry and the levels in right SN relative to the left SN were determined for each mouse. There was no significant difference in expression between males and females, thus, the data was combined for subsequent analysis. The results showed that the LV/dSaCas9-DNMT3A vector significantly repressed total human α-syn expression in the right SN compared to the left SN (-23.9% ± 12, t(9)=2.72 p=0.014, paired t-test, Fig. 3c-d). There were no significant differences in the expression of α-syn pS129 or aggregated α-syn. To evaluate the impact on dopaminergic neurons we assessed endogenous mouse TH expression in the SN and the striatum. There was no significant difference in mouse TH expression in the SN between the right and left hemisphere (Fig. 3e), however, there was significantly higher expression of mouse TH in the right striatum (+52% ± 17, t(9)=3.76 p=0.0045, paired t-test, Fig. 3c-d), indicating a greater retention of dopaminergic neuron projections in the presence of the LV/dSaCas9-DNMT3A. While these results support our previous *in vitro* findings, showing that LV/dSaCas9-DNMT3A can downregulate α-syn expression, the effects were not sufficient to significantly impact pathologically relevant hallmarks of PD, *i.e*. α-syn pS129 and aggregated α-syn.

**Figure 3.**
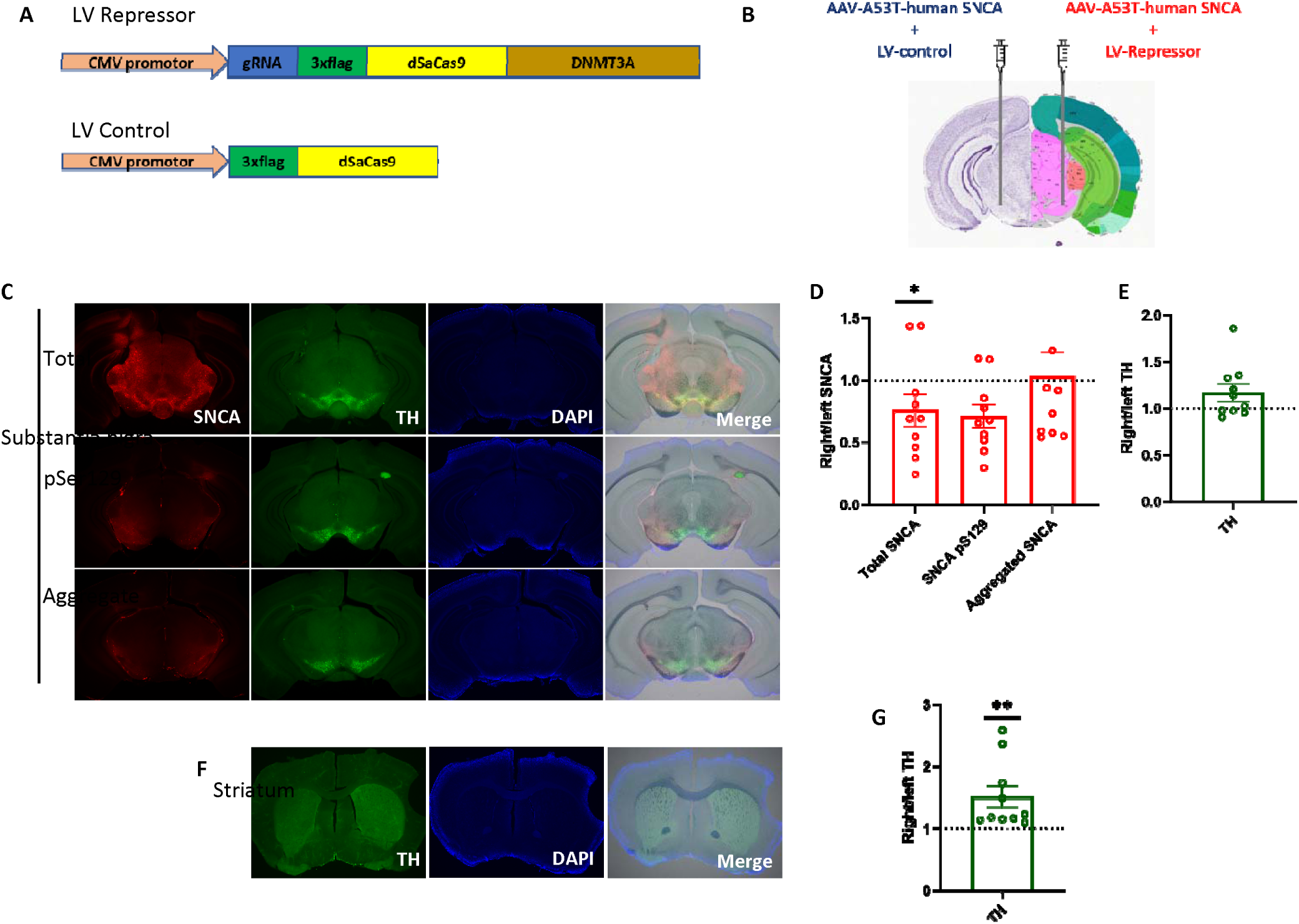
LV/dSaCas9-DNMT3A reduces. α**-synuclein expression in PD mouse model** (**A**) Schematic showing the structure of the LV/dSaCas9-DNMT3A vector. (**B**) Diagram of experimental design; bilateral stereotaxic injections. AAV/A53T-hSNCA was co-injected with LV/dSaCas9-DNMT3A into the right SN of mice, and AAV/A53T-hSNCA was co-injected with the LV/dCas control (no repressor, no gRNA) into the left SN. Image modified from the Allen Brain Atlas (allenbrainatlas.org). (**C**) Representative images of brain coronal slices at 2x magnification 6 weeks post-injection, showing protein expression of human total α-syn, α-syn pS129, and aggregated α-syn, and mouse TH, and DAPI staining in the SN. Fluorescence intensity was quantified using ImageJ. Box plot displays the ratios of the right SN relative to the left SN. Each open circle represents the quantified signal right/left for a mouse. (**D)** The LV/dSaCas9-DNMT3A vector significantly repressed total human α-syn expression in the right SN compared to the left SN. There were no significant differences in the expression of α-syn pS129 or aggregated α-syn. (**E**) There was no significant difference in mouse TH expression between the right and left SN. (**F**) Representative images of brain coronal slices at 2x magnification 6 weeks post-injection, showing protein expression of mouse TH, and DAPI staining in the striatum. (**G**) Expression of mouse TH was significantly higher in the right striatum in the presence of the LV/dSaCas9-DNMT3A vector. Values represent mean ratios ± SEM. N =10 (6 female, 4 male) * p < 0.05 ** p < 0.01; Paired t-test.

### LV/dSaCas9-KRAB-MeCP2(TRD) reduces pathologic forms of α-synuclein in PD mouse model

We recently demonstrated that d*Sa*Cas9-KRAB-MeCP2(TRD) vector platform represses robustly and sustainably the expression of multiple targets, *in vitro* and *in vivo* ^63^. LV/dSaCas9-KRAB-MeCP2(TRD) displayed substantially enhanced repression capacity *in vitro* compared to dSaCas9-DNMT3A (Fig. 1). In addition, expression of the KRAB-MeCP2 synthetic repressor molecule vectors driven by dopaminergic and cholinergic specific regulatory sequences resulted in robust and sustained neuronal-type specific reduction of α-syn and amelioration of pathological cellular phenotypes ^67^. To investigate the *in vivo* efficacy of the LV/dSaCas9-KRAB-MeCP2(TRD) (Fig. 4a) system, a total of ten C57B/L6 mice (5 males, 5 females) received bilateral stereotaxic injections. AAV/hA53T-*SNCA* was co-injected with LV/dSaCas9-KRAB-MeCP2(TRD) (7.5×10^6^ vg) into the right SN of mice, and AAV/hA53T-*SNCA* (2.5×10^9^ vg) was co-injected with the LV/dSaCas9 control (no repressor, no gRNA; 7.5×10^6^ vg) into the left SN (Fig. 4b). At six weeks post-injection, the expression of total human α-syn, α-syn pS129, and aggregated α-syn was quantified by immunohistochemistry and the levels in right SN relative to the left SN were determined for each mouse. There was no significant difference in expression between males and females, thus, the data was combined for subsequent analysis. The LV/dSaCas9-KRAB-MeCP2(TRD) vector significantly repressed total human α-syn expression in the right SN compared to the left SN (-40% ± 10, t(9)=3.4 p=0.0074, paired t-test, Fig. 4c-d). Expression of α-syn pS129 and aggregated α-syn were repressed by a even greater magnitude (pS129: -70.84% ± 7.9, t(9)=5.25 p=0.0005. Aggregated: -72.59% ± 6.4, t(9)=6.66 p=0.0001, paired t-tests, Fig. 4c-d). We also assessed endogenous mouse TH expression in the SN and the striatum. In both the SN (Fig. 4c, e) and the striatum (Fig. 4f-g) TH expression was significantly higher in the presence of the LV/dSaCas9-KRAB-MeCP2(TRD) vector (SN: +1.2fold, t(9)=2.27 p=0.049. Striatum: +2.6-fold, t(9)=4.11 p=0.0026, paired t-test), indicating a greater retention of dopaminergic neurons and projections.

**Figure 4.**
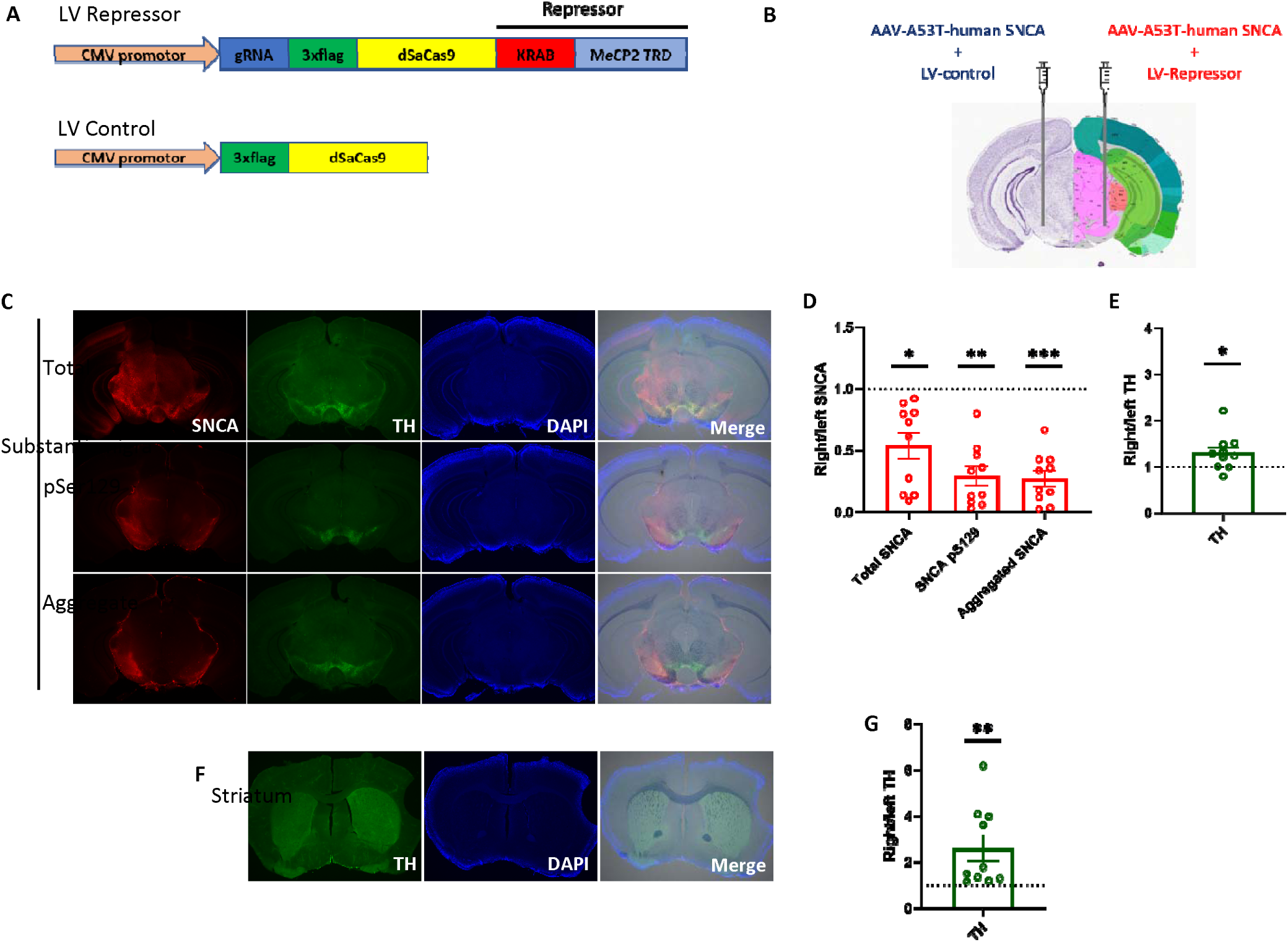
LV/dSaCas9-KRAB-MeCP2(TRD) reduces pathologic forms of α-synuclein. (**A**) Schematic showing the structure of the LV/dSaCas9-KRAB-MeCP2(TRD) vector. (**B**) Diagram of experimental design; bilateral stereotaxic injections. AAV/A53T-hSNCA was co-injected with LV/dSaCas9-KRAB-MeCP2(TRD) into the right SN of mice, and AAV/A53T-hSNCA was co-injected with the LV/dCas control (no repressor, no gRNA) into the left SN. Image modified from the Allen Brain Atlas (allenbrainatlas.org). (**C**) Representative images of brain coronal slices at 2x magnification 6 weeks post-injection, showing protein expression of human total α-syn, α-syn pS129, and aggregated α-syn, and mouse TH, and DAPI staining in the SN. Fluorescence intensity was quantified using ImageJ. Box plot displays the ratios of the right SN relative to the left SN. Each open circle represents the quantified signal right/left for a mouse. (**D)** The LV/dSaCas9-KRAB-MeCP2(TRD) vector significantly repressed expression of total human α-syn, α-syn pS129 or aggregated α-syn in the right SN compared to the left SN. (**E**) Expression of mouse TH was significantly higher in the right striatum in the presence of the LV/dSaCas9-KRAB-MeCP2(TRD) vector. (**F**) Representative images of brain coronal slices at 2x magnification 6 weeks post-injection, showing protein expression of mouse TH, and DAPI staining in the striatum. (**G**) Expression of mouse TH was significantly higher in the right striatum in the presence of the LV/dSaCas9-KRAB-MeCP2(TRD) vector. Values represent mean ratios ± SEM. N =10 (5 female, 5 male) * p < 0.05 ** p < 0.01 *** p < 0.001; Paired t-test.

**Figure 5.**
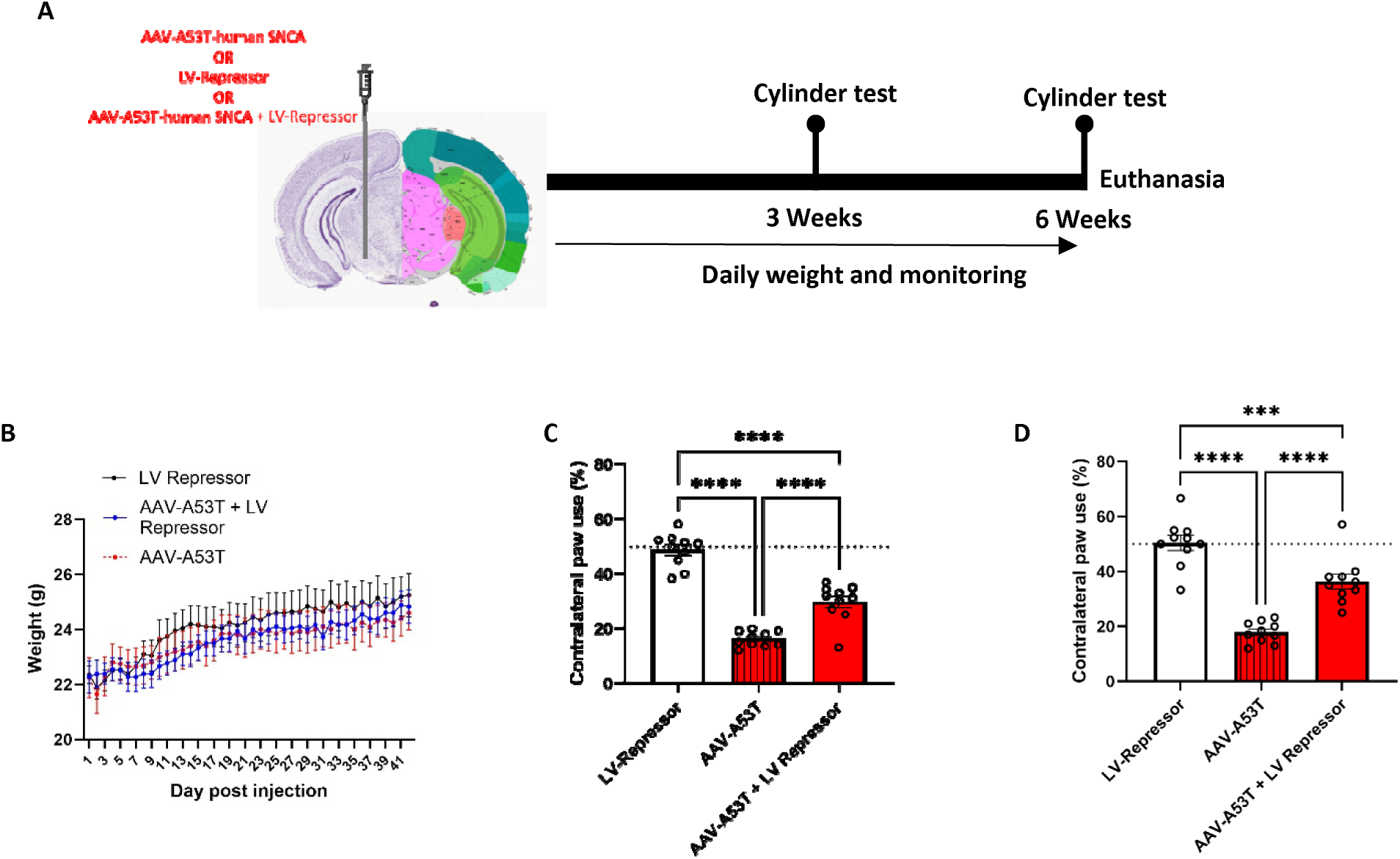
LV/dSaCas9-KRAB-MeCP2(TRD) rescues motor impairment in PD mouse model. (**A**) Diagram of experimental design; unilateral stereotaxic injections into the SN of either: LV/dSaCas9-KRAB-MeCP2, AAV/A53T-hSNCA, or a co-injection of AAV/A53T-hSNCA + LV/dSaCas9-KRAB-MeCP2. Image modified from the Allen Brain Atlas (allenbrainatlas.org). Cylinder tests were performed at 3 weeks and 6 weeks post-injection, animal weights were monitored throughout the experiment. (**B**) Animal weights increased throughout the duration of experiment and did not significantly differ between groups (**C**) At 3 weeks, unilateral AAV/A53T-hSNCA delivery leads to a significant asymmetry of forepaw use with preference of the ipsilateral forepaw relative to the side of injection as compared to LV/dSaCas9-KRAB-MeCP2 (alone) control mice. LV/dSaCas9-KRAB-MeCP2 co-administration with AAV/A53T-hSNCA significantly improved contralateral paw use, though not to the level of control (LV/dSaCas9-KRAB-MeCP2 alone) mice. (**D**) At 6 weeks, the contralateral forepaw use deficit induced by AAV/A53T-hSNCA delivery persists, the AAV/A53T-hSNCA + LV/dSaCas9-KRAB-MeCP group showed increased improvement in the use of the impaired limb. Values represent mean ± SEM. N =12 females per group *** p < 0.001 **** p < 0.0001; One-way ANOVA followed by Tukey test.

### LV/dSaCas9-KRAB-MeCP2(TRD) rescues motor impairment in PD mouse model

To explore the behavioral consequences of the α-syn repression and dopaminergic neuron retention, we assessed motor deficits by the cylinder test at 3 and 6 weeks timepoints. C57B/L6 mice (10 female per group) received unilateral injection into the SN of (1) LV/dSaCas9-KRAB-MeCP2(TRD), or (2) AAV/hA53T-*SNCA*, or (3) a co-injection of AAV/hA53T-*SNCA* + LV/dSaCas9-KRAB-MeCP2(TRD) (Fig. 5a). Animal weights and welfare were monitored over the 6-week duration of the experiment, weights increased throughout the experiment and did not significantly differ between groups (F(2,26) = 0.25, p = 0.78, two-way ANOVA, Fig. 5b). A significant asymmetry of forepaw use, with preference for the side ipsilateral to injection was found in both the AAV/hA53T-*SNCA* group (16.23% ± 0.8), and AAV/hA53T-*SNCA* + LV/dSaCas9-KRAB-MeCP2(TRD) group (29.78% ± 2.1) at 3 weeks, compared to the LV/dSaCas9-KRAB-MeCP2(TRD) group where no deficit was observed (48.75% ± 1.8, Fig. 5c). Notably, the motor deficit was significantly ameliorated in the AAV/hA53T-*SNCA* + LV/dSaCas9-KRAB-MeCP(TRD) group compared to the AAV/hA53T-*SNCA*, demonstrating that our LV-Repressor system successfully ameliorated motor deficit induced in our PD mouse model (F(2,27) = 86.29, p<0.0001, one-way ANOVA, Fig. 5c). Tukey’s HSD Test for multiple comparisons confirms that the mean value for contralateral forepaw use was significantly different between: LV/dSaCas9-KRAB-MeCP2(TRD) and AAV/hA53T-*SNCA* (p < 0.0001, 95% C.I. = [26.36, 38.69]), AAV/hA53T-*SNCA* and AAV/hA53T-*SNCA* + LV/dSaCas9-KRAB-MeCP2(TRD) (p < 0.0001, 95% C.I. = [12.8, 25.14]), and, LV/dSaCas9-KRAB-MeCP2(TRD) and AAV/hA53T-*SNCA* + LV/dSaCas9-KRAB-MeCP2(TRD). (p < 0.0001, 95% C.I. = [-19.72, -7.39], Fig. 5c). Importantly, and consistently with our recent results ^63^ these differences were maintained at 6 weeks, demonstrating the durability of the effect on behavioral assessment. No motor impairment of the contralateral paw was observed in LV/dSaCas9-KRAB-MeCP2(TRD) group (50.39% ± 2.7), and in the AAV/hA53T-*SNCA* group impairment was maintained (17.88% ± 1.2). However, the AAV/hA53T-*SNCA* + LV/dSaCas9-KRAB-MeCP(TRD) group showed increased improvement in the use of the impaired limb (34.88% ± 2.7, F(2,27) = 45.57, p<0.0001, one-way ANOVA, Fig. 5d). Tukey’s HSD Test for multiple comparisons confirms that the mean value for contralateral forepaw use was significantly different between: LV/dSaCas9-KRAB-MeCP2(TRD) vs AAV/hA53T-*SNCA* (p < 0.0001, 95% C.I. = [24.07, 40.96]), AAV/hA53T-*SNCA* vs AAV/hA53T-*SNCA* + LV/dSaCas9-KRAB-MeCP2(TRD) (p = 0.0003, 95% C.I. = [7.06, 23.96]), and, LV/dSaCas9-KRAB-MeCP2(TRD) vs AAV/hA53T-*SNCA* + LV/dSaCas9-KRAB-MeCP2(TRD). (p < 0.0001, 95% C.I. = [-25.45, - 8.56], Fig. 5d).

Overall, these results indicated that our LV/dSaCas9-KRAB-MeCP2(TRD) epigenomic therapy platform can effectively and sustainably reduce pathological forms of α-syn, resulting in the retention of dopaminergic neurons and rescue of motor impairment, and by that displaying the potential as a therapeutic strategy for PD.

### Safety profile of LV/dSaCas9-KRAB-MeCP2(TRD)

Given the promising results on the efficacy of the LV/dSaCas9-KRAB-MeCP2(TRD) platform, we conducted pilot safety evaluations including daily weights, serum chemistry and complete blood count (CBC) analysis of mice treated with the repressor. C57B/L6 mice received unilateral injection into the SN of either: LV/dSaCas9-KRAB-MeCP2(TRD), LV/dSaCas9 control, or saline (Fig. 6a). Male: LV/dSaCas9-KRAB-MeCP2(TRD) n=4, LV/dSaCas9 control n=6, or saline n=9. Female: LV/dSaCas9-KRAB-MeCP2(TRD) n=11, LV/dSaCas9 control n=6, or saline n=11. Animal weights and welfare were monitored over the 9-week duration of the experiment, both male and female weights increased throughout the experiment and did not significantly differ between groups (Male: F(2, 17) = 0.5740, p = 0.57, Female: F(2, 15) = 1.391, p = 0.28, two-way ANOVA, Fig. 6b-c). CBC analysis revealed: in females there was no significant effect of treatment on white blood cell count (WBC), in males, a lower WBC count was observed with both the LV-Repressor and LV-Control when compared to the saline (F(2, 41) = 11.26, p=0.0001, two-way ANOVA; Tukey’s HSD test: Male Saline vs LV/dSaCas9-KRAB-MeCP2(TRD) p = 0.0002, 95% C.I. = [0.53, 1.9], Male Saline vs LV/dCas control p = 0.0003, 95% C.I. = [0.45, 1.6], Fig. 6d). There was no significant effect of treatment on red blood cell (RBC) or platelet count in either males or females (Fig. 6e-f).

**Figure 6.**
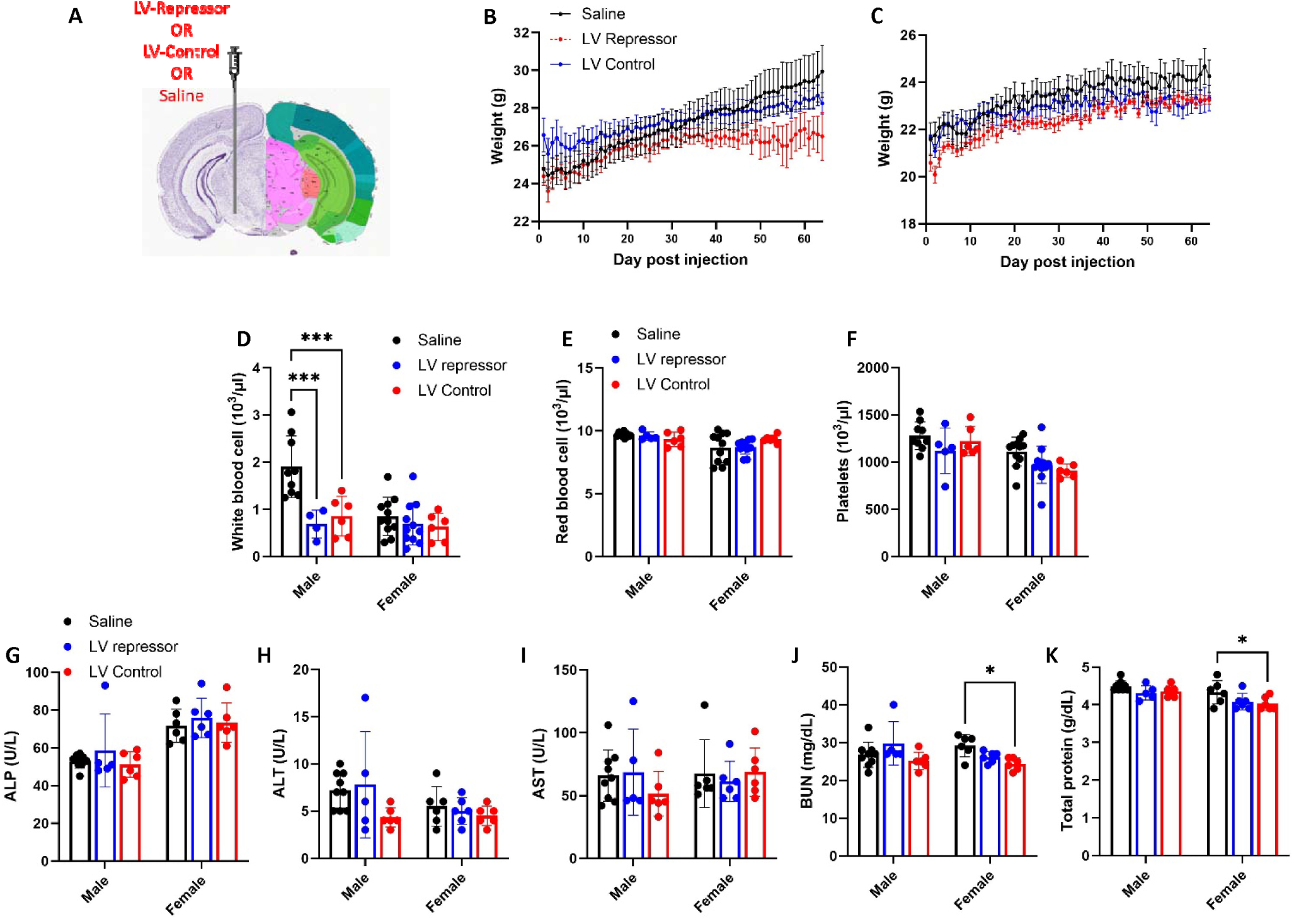
Safety profile of LV/dSaCas9-KRAB-MeCP2(TRD) (**A**) Diagram of experimental design; unilateral stereotaxic injections into the SN of either: LV/dSaCas9-KRAB-MeCP2, LV/dCas control, or saline. Image modified from the Allen Brain Atlas (allenbrainatlas.org). Animal weights were monitored over the 9 week duration of the experiment, both (**B**) male and (**C**) female weights increased throughout the experiment and did not significantly differ between groups. CBC analysis: (**D**) In males, there was a lower WBC count with both the LV-Repressor and LV-Control when compared to the saline. In females, there were no differences in WBC count between groups. In both males and females there were no differences in (**E**) RBC count or (**F**) Platelets between groups. Serum chemistry: In both males and females there were no differences in (**G**) ALP (**H**) ALT (**I**) AST between groups. In females, slightly lower levels of both(**J**) BUN and (**K**) total protein were observed with the LV-Control when compared to the saline. In males there were no differences between groups. Values represent mean ± SEM. Male: LV/dSaCas9-KRAB-MeCP2 n=4, LV/dCas control n=6, or saline n=9. Female: LV/dSaCas9-KRAB-MeCP2 n=11, LV/dCas control n=6, or saline n=11 * p < 0.05 *** p < 0.001; Two-way ANOVA followed by Tukey test.

Serum chemistry was analyzed to assess liver and kidney function. Alkaline phosphatase (ALP) Alanine aminotransferase (ALT), and aspartate aminotransferase (AST) are enzymes that are primarily found in the liver and are important indicators of liver health and function. Neither LV/dSaCas9-KRAB-MeCP2(TRD) injection or LV/dSaCas9 control altered the expression of ALP, ALT or AST in either males or females when compared to saline (Fig. 6g-i). Blood urea nitrogen (BUN) and total protein were analyzed as a measure of kidney function. In males there was no significant effect of treatment on BUN or total protein. In females, slightly lower levels of both BUN and total protein were observed with the LV-Control when compared to the saline (BUN: F(2, 32) = 4.317, p=0.022, two-way ANOVA; Tukey’s HSD test: Female Saline vs LV/dSaCas9 control p = 0.035, 95% C.I. = [0.28, 9.4]. Total protein: F(2, 32) = 5.29, p=0.01, two-way ANOVA; Tukey’s HSD test: Female Saline vs LV/dCas control p = 0.04, 95% C.I. = [0.007, 0.59] Fig. 6j-k). CBC and serum chemistry data as means ± SEM are provided in Table S1. Overall, these analyses did not reveal any major signs of toxicity or safety issues after the stereotaxic injection of LV/dSaCas9-KRAB-MeCP2(TRD) system for α-syn repression mediated by epigenome editing.

## DISCUSSION

The current study is the first *in vivo* validation of our novel *SNCA*-targeted epigenome-editing technology. The preclinical experiments here demonstrate the therapeutic potential of our technology for PD and related synucleinopathies. We developed two *all-in-one* LV vectors harboring dSaCas9, gRNA targeted at *SNCA*-intron 1, and one of the two repressor molecules (1) DNMT3A or (2) engineered fusion of KRAB and MeCP2(TRD). Administration of both vectors into the SN of a novel PD mouse model resulted in a precise and efficient downregulation of the human α-syn protein. However, our data showed that the synthetic KRAB-MeCP2(TRD) molecule displayed enhanced repressor capabilities in both *in vitro* and *in vivo* models.

Moreover, α-syn reduction driven by the KRAB-MeCP2(TRD) was sufficient to rescue PD-associated pathological and behavioral characteristics of our PD mouse model including, lowering the levels of pS129 and aggregated α-syn protein, retaining dopaminergic neurons, and ameliorating motor impairment. By that, this pre-clinical study advanced our previous work towards investigational new drug (IND) enablement.

Previously, we edited the DNA methylation and chromatin accessibility landscape within *SNCA* intron 1 sites in hiPSC-derived dopaminergic and cholinergic neurons from a PD/DLB patient and demonstrated target engagement and the beneficial effects on neuropathological and cellular phenotypes ^59, 60^. While human relevant models such as patient hiPSC-derived neuronal models are valuable tools in the early stages of drug discovery, several aspects in drug developments such as, behavioral and safety assessments, require the whole organism level. These constrains are even more pronounced for CNS disorders due to the complex anatomical and functional organization of the brain. Thus, the progression into *in vivo* models is necessary for advancement towards IND enablement and clinical trials. To tackle these limitations, we developed a new humanized PD-mouse model induced by overexpression of hA53T α-syn protein. PD models based on AAV driven overexpression of hA53T α-syn in the SN are well-established, characterized by α-syn protein aggregation, degeneration of nigral dopaminergic neurons, decreases in TH, and motor deficits in mice, rats and monkeys ^66, 68, 69^. Leveraging on these previous AAV/hA53T-*SNCA* based models, our transgene is expressed via human native regulatory sequences derived from the promoter/intron 1 region to imitate the expression pattern observed in humans. We recapitulated the pathological and behavioral characteristics described previously ^66, 68, 69^.

In this work, we further refined the therapeutic technology for finetuning the production of α-syn. Foremost, we reduced the overall size of the *all-in-one* transgene to improve the transduction efficiently and adaptability for other packaging methods. First, the new system comprises the smaller dSaCas9. Second, we engineered a synthetic repressor molecule, a fusion of MeCP2’s transcription repression domain (TRD) and KRAB. The KRAB-MeCP2(TRD) molecule is smaller in size than MeCP2 as well as DNMT3A. Furthermore, we recently demonstrated that the developed KRAB-MeCP2(TRD) system is more efficient and sustainable compared with the KRAB, or MeCP2 counterparts for the targeted repression of the genes, in vitro and in vivo ^63^. These modifications have improved the effectiveness of our epigenome-editing system and enhanced adaptability for packaging into alternative viral vectors such as AAV capsids. Additionally, the prospects for a one-time treatment that could permanently reduce expression of pathologically overexpressed genes is poised to bring this gene therapy system to wide-range of neurodegenerative and other diseases ^63^.

A number of studies have reported the advantageous effects of reducing α-syn expression using RNA based technologies that target *SNCA*-mRNA. Antisense oligonucleotides (ASOs) were shown to reduce production of α-syn, leading to prevention α-syn pathology and dopaminergic cell dysfunction in rodent preformed fibril (PFF) models of PD ^46^. shRNA mediated knockdown of α-syn has been reported to be neuroprotective in both SNCA-Tri NPCs, and in a rodent rotenone model of PD ^47, 70^. Both ASOs and RNAi have also been successfully employed to reduce α-syn production in non-human primates ^46, 71^. However, these strategies have resulted in a nigrostriatal degeneration as a direct outcome of α-syn knockdown. AAV expressing shRNA or siRNA targeting endogenous α-syn, initiated robust reduction of α-syn resulting in a rapid loss of nigrostriatal dopaminergic neurons and striatal dopamine in rodents and non-human primates ^48–50^. The results suggested that the divergence between these findings is due to differences in the degree of α-syn knockdown, that is to say, neuroprotective effects were reported when the level of α-syn repression reached to 35-55%, while a more robust knockdown led to toxicity ^47, 71^. It has been suggested ^48^ that there is a ∼50% critical threshold of α-syn repression, below which a pathological cascade is imitated leading to dopaminergic neuron loss. Patients with triplication or duplication of the *SNCA* locus showed 200% and 150% higher levels of *SNCA* production, respectively, and manifest proportionally early-onset PD symptoms ^37, 38^. Consequently, an estimated therapeutic window of 30-50% reduction in α-syn expression should find balance between therapeutic value and restoration of normal endogenous α-syn levels. Here, we achieved a reduction in α-syn expression with both the LV/dSaCas9-DNMT3A vector and the LV/dSaCas9-KRAB-MeCP2(TRD) vectors. However, the magnitude of repression with the later vector was greater (23.9% vs 40%). The 40% reduction accomplished with the LV/dSaCas9-KRAB-MeCP2(TRD) system meets the mid-point of the theorized therapeutic window and coincided with reduction in α-syn pS129 and α-syn aggregates – hallmarks of PD neuropathology. Indeed, this reduction was sufficient to ameliorate the retention of dopaminergic neurons in the SN and striatum, and to improve motor deficits characteristics of our PD model. On the other end, the vector expressing the DNMT3A repressor did not reach this therapeutic window for α-syn repression (23.9%), which may explain the undetectable effects on PD associated neuropathological features. This highlights the advancement of our *SNCA*-targeted epigenome therapy development carrying the synthetic KRAB-MeCP2(TRD) molecule as a therapeutic for PD and other synucleinopathies.

Overexpression of α-syn disrupts dopaminergic neurotransmission leading to motor impairments. The onset of motor symptoms in PD patients is preceded by nigrostriatal degeneration with an estimated ∼30% loss of SN neurons, and an even higher loss of striatal terminals ^72^. Similarly, the AAV/hA53T-*SNCA* mouse models showed significant degeneration of dopaminergic neurons in the SN and a significant loss of nigrostriatal fibers in the striatum ^66, 68, 69, 73^. In agreement with these evidences, we observed a significant decrease in TH in the striatum of the AAV/hA53T-*SNCA* SN injected model. The LV/dSaCas9-KRAB-MeCP2(TRD) vector significantly increases TH expression in both the SN and the striatum, suggesting a retention of dopaminergic neurons. This is supported by our behavioral data, showing the reversal of motor impairment in our AAV/hA53T-*SNCA* treated mice as early as 3 weeks and sustainable through 6 weeks post injection. Overall, the results supported the efficiency and sustainability of the *SNCA* gene repression.

There are a few limitations and next steps to consider as we work to develop this therapeutic technology for clinical application. First, this LV/dSaCas9-KRAB-MeCP2(TRD) vector is expressed from a constitutive promotor (CMV) which carries the risk of undesirable manipulation of SNCA in cell types that are not vulnerable to PD. We have recently modified the system to include either TH or choline acetyltransferase (ChAT) promoters that are specifically expressed in dopaminergic and cholinergic neurons, respectively ^67^. Validation studies in hiPSC-derived mature neurons demonstrated neuronal-type specificity and suppression of α-syn resulting in rescuing disease related pathological hallmarks of synucleinopathies in the vulnerable neuronal types for PD and DLB ^67^. This strategy targets cell-type specific expression of the therapeutic vector and hence reduces the risk of unwanted effects on other cell types, and opens the opportunity to explore other routes of administration (ROA) such as intracerebroventricular delivery, that may be more feasible in a clinical setting.

Secondly, LV vectors were used in the development of this system due to their large payload-capacity and ability to sustain robust gene expression. Our pilot safety assessment of the LV/dSaCas9-KRAB-MeCP2(TRD) system did not raise any immediate safety concerns. However, concerns have been raised related to the ability of LV to integrate into the host genome, that could result in unintended off-target effects, long-term expression issues, and an increased risk of insertional mutagenesis and oncogenicity ^74, 75^. AAV is a non-pathogenic, transiently expressing system, with very low integration capacity, and can sustain long-term transgene expression. AAV vectors have emerged as the preferred choice in clinical applications including for CNS disorders ^76^. We have demonstrated that the d*Sa*Cas9-KRAB-MeCP2(TRD) system can be successfully packaged into an AAV vector ^63^, effectively and sustainably repressing the expression of multiple genes of interest, both *in vitro* and in vivo, including ApoE, the strongest genetic risk factor for late-onset Alzheimer’s disease ^63^. Next steps to move towards clinical application will include combining these modifications to develop a dopaminergic-neuron specific AAV packaged system for safe, efficient and precise delivery of the SNCA-targeted epigenome-editing system. Furthermore, extension of our technology development to treat other synucleinopathies, via the development of other cell-type specific regulatory elements to drive the specific expression of the therapeutic vector in particular cell types affected in each disease within the broader spectrum of synucleinopathies.

In conclusion, we demonstrated the efficacy of *SNCA*-targeted epigenome therapy in a PD mouse model, effectively repressing *SNCA* expression to a level that reverses PD-associated neuropathological hallmarks and behavioral motor impairments. This study represents an advancement towards precision gene-targeted therapy for PD and other synucleinopathies.

## MATERIALS AND METHODS

### Plasmid design and construction

To create SaCas9-based constructs, we used pX603-AAV-CMV:NLS-dSaCas9(D10A,N580A)-NLS-3xHA-bGHpA that was obtained from addgene (plasmid #61594; gift from Fang Zhang’s laboratory). The amplified fragment of 3202bp harboring dCas9 CDS was cloned into pBK694 as described above. The primers contained AgeI-BamHI sites which were used for cloning, as above. The resulting plasmid was pBK1124. The oligo flanked by NdeI-NotI sites was annealed and cloned to create the pBK1129 plasmid. We then amplified the C-terminus of *Sa*Cas9 and replaced it with the mutated version that carried no stop codon. The following oligo was used 5’-ggatcctcaaataaaagatctttgttttcattagatctgtgtgttggttttttgtgtgcggccggtacc-3’ to remove the stop codon at the C-terminus of *Sa*Cas9. This plasmid was named pBK1198. Then, we introduced the adaptor sequence downstream from C-terminus of *Sa*Cas9. The sequence was 5’-GATCCggtggaggaagtggcgggtcagggtcgggtggcACTAGTataGCTAGCggaggtggttcgccaaagaagaaacggaaggtgG-3’. The resulting plasmid is pBK1294b. This plasmid has been used as the intermediate vector for cloning all of the d*Sa*Cas9-repressor fragments, including DNMT3A and KRAB-MeCP2(TRD).

The following gRNA oligos targeting the CAG/Intron1 promoter were selected: gRNA1 5’-gacctcccacacctggccca-3’; gRNA2 5’-gctcagggtagatagctgagg-3’. We derived the CAG promoter from pBK814-pLenti-CAG-GFP-p2a-Nanoluc-WPRE. The promoter was cloned into pBK533 to create pLV-CAG -GFP-P2A-Nluc-WPRE. The resulting vector pBK573 has been recorded. The SNCA intron 1-alpha syn ORF has been synthesized using GenScript’s DNA synthesis service. The oligo was flanked with the ClaI-AgeI restriction sites used to clone it into the above construct to create pCL35 plasmid. Plasmid sequences are provided in Figure S1.

### Vector Production

Lentiviral vectors were generated using the transient transfection protocol, as described previously ^77^. Briefly, 15 μg vector plasmid, 10 μg psPAX2 packaging plasmid (Addgene 12260 generated in Dr. Didier Trono’s lab, EPFL, Switzerland), 5 μg pMD2.G envelope plasmid (Addgene 12259, generated in Dr. Trono’s lab), and 2.5 μg pRSV-Rev plasmid (Addgene 12253, generated in Dr. Trono’s lab) were transfected into 293T cells. Vector particles were collected from filtered conditioned medium at 72 hr post-transfection. The particles were purified using the sucrose-gradient method ^77^, and concentrated >250-fold by ultracentrifugation (2 hours at 20,000 rpm). Vector and viral stocks were aliquoted and stored at −80°C.

### Titer of Vector Preparations

Titers were determined for the vectors expressing puromycin selection marker by counting puromycin-resistant colonies as described ^77^ and by p24^gag^ ELISA method equating 1 ng p24gag to 1 × 104 viral particles. The MOI was calculated by the ratio of the number of viral particles to the number of cells. The p24^gag^ELISA was carried out as per the instructions in the HIV-1 p24 antigen capture assay kit (NIH AIDS Vaccine Program). Briefly, high-binding 96-well plates (Costar) were coated with 100 μL monoclonal anti-p24 antibody (NIH AIDS Research and Reference Reagent Program, catalog 3537) diluted 1:1,500 in PBS. Coated plates were incubated at 4°C overnight, then blocked with 200 μL 1% BSA in PBS and washed three times with 200 μL 0.05% Tween 20 in cold PBS. Next, plates were incubated with 200 μL samples, inactivated by 1% Triton X-100 for 1 hr at 37°C. HIV-1 standards (catalog SP968F) were subjected to a 2-fold serial dilution and applied to the plates at a starting concentration equal to 4 ng/mL. Samples were diluted in RPMI 1640 supplemented with 0.2% Tween 20 and 1% BSA, applied to the plate and incubated at 4°C overnight. Plates were then washed six times and incubated at 37°C for 2 hr with 100 μL polyclonal rabbit anti-p24 antibody (catalog SP451T), diluted 1:500 in RPMI 1640, 10% fetal bovine serum (FBS), 0.25% BSA, and 2% normal mouse serum (NMS; Equitech-Bio). Plates were then washed as above and incubated at 37°C for 1 hr with goat anti-rabbit horseradish peroxidase immunoglobulin G (IgG) (Santa Cruz Biotechnology), diluted 1:10,000 in RPMI 1640 supplemented with 5% normal goat serum (NGS; Sigma), 2% NMS, 0.25% BSA, and 0.01% Tween 20. Plates were washed as above and incubated with 3,3’,5,5’-tetramethylbenzidine (TMB) peroxidase substrate (KPL) at room temperature for 10 min. The reaction was stopped by adding 100 μL 1N HCL. Plates were read by Microplate Reader (The iMark Microplate Absorbance Reader, Bio-Rad) at 450 nm and analyzed in Excel. All experiments were performed in triplicates.

### Cell culture

The HEK293T/pCLl35 cell line was generated through transduction of HEK293T cells (ATCC® CRL3216™) with pLenti-pCL35 and selected using puromycin. These cells express Alpha-syn ORF attached to dGFP and luciferase (NLuc) downstream of a CAG/SNCA Intron 1 promoter. The dGFP tag provides visual changes of gene repression whereas the luciferase provides higher spatial resolution identifying protein concentration changes over time. Maintenance cells were grown in DMEM (Gibco), supplemented with 10% fetal bovine serum (Gibco), penicillin/streptomycin 1% (Thermo Fisher Scientific), 2 mM L-glutamine, 1% MEM NEAA (Gibco), and 1 mM sodium pyruvate (Gibco). To note: the cells were transduced with the MOI = 0.1 to ensure 1 copy/cell following the selection. Before lentiviral vector transduction, the cells were seeded at 2.5 × 105 cells per 12-well plate and cultured in DMEM with 2% FBS with no supplements. HEPG2 cells were maintained in DMEM (Gibco), supplemented with 10% fetal bovine serum (Gibco), penicillin/streptomycin 1% (Thermo Fisher Scientific), 2 mM L-glutamine, 1% MEM NEAA (Gibco), and 1 mM sodium pyruvate (Gibco).

### Vector transduction

At 50% confluency, the HEK293T/pCL35 line was transduced with lV/dCas9-repressor vectors. The following vectors were used: LV/dSaCas9-KRAB-MeCP2(TRD) and LV/dSaCas9 - DNMT3A vectors. The vectors were used with two different gRNAs (gRNA1 and gRNA2) (see above). The vectors were used at the MOIs = 2 vg/cell. The cells were harvested at day 4 post-transduction (p.t.).

### Luciferase reporter assay

Cells from each 12-well plate were first harvested and washed twice with 1× PBS before being resuspended in 200 µL of 1× PBS. 50 µL of the cell and 1× PBS mixture were transferred into a 96-well plate bottom white plate (Costar Cat#3922) and lysis buffer from Nano-Glo Luciferase Assay Kit (Promega Cat#N1120) was added directly to the plates following the manufacturer’s protocol. The data was obtained using a microplate spectrophotometer (Bio-Rad, Hercules, CA). Total protein concentration, which was determined by the DC Protein Assay Set (Bio-Rad, Hercules, CA), was then used for data normalization.

### Stereotaxic injections

All experiments involving animal use were performed in accordance with the ethical guidelines of Duke Institutional Animal Care and Use Committee. Male and female C57BL/6 mice weighing 20–30g were obtained from Charles River Laboratories and investigated at the age of 16 weeks. Mice were kept under standard conditions (21 °C, 12 h/12 h light-dark cycle) with food and water available *ad libitum* in their home cages. Animal welfare and weights were monitored daily for 42 days.

Under isoflurane anesthesia (0.5–2% isoflurane in O2) mice were injected either unilaterally or bilaterally (as depicted in each Fig) into the SN via a Neuros syringe (Hamilton) at a rate of 150 nl/minute using the following stereotaxic coordinates (relative to bregma): -3 mm anterior, ± 1.4 mm lateral, and 4.55 mm ventral.

Model validation (Fig 2): Bilateral injections: AAV/hA53T-*SNCA* (1 µl) into one hemisphere, AAV/null (1 µl) control into other hemisphere.

LV/dSaCas9-DNMT3A (Fig 3): Bilateral injections: AAV/hA53T-*SNCA* and LV/dSaCas9-DNMT3A combined (1 µl) into one hemisphere, AAV/hA53T-*SNCA* and LV/dSaCas9-Control combined (1 µl) into other hemisphere.

LV/dSaCas9-KRAB-MeCP2(TRD) (Fig 4): Bilateral injections: AAV/hA53T-*SNCA* and LV/dSaCas9-KRAB-MeCP2(TRD) combined (1 µl) into one hemisphere, AAV/hA53T-*SNCA* and LV/dSaCas9-Control combined (1 µl) into other hemisphere.

Behavior (Fig 5): Unilateral injection of either AAV/hA53T-*SNCA* (1 µl), LV/dSaCas9-KRAB-MeCP2(TRD) (1 µl), or AAV/hA53T-*SNCA* and LV/dSaCas9-KRAB-MeCP2(TRD) combined (1 µl), into one hemisphere.

Safety (Fig 6): Unilateral injection of either LV/dSaCas9-KRAB-MeCP2(TRD) (1 µl), or Saline (1 µl), into one hemisphere.

### Immunohistochemistry and Imaging

At 42 days post-injection, mice were transcardially perfused with 10% formalin and coronal slices (100 μm) containing the SN and striatum were cut with a vibratome (VT1000S, Leica Microsystems). For immunohistochemistry, brain slices were immersed for 1 h at room temperature in PBS containing 0.2% Triton-X 100, 5% normal goat serum and 1x fish gelatin, then incubated with primary antibody for total human α-syn (1:1000, ab138501, Abcam), α-syn pS129 (1:1000, ab184674, Abcam), or aggregated α-syn (1:1000, MABN389, MilliporeSigma), and mouse TH (1:1000, ab76442, Abcam) at 4°C overnight on a rotator. The next day, the slices were rinsed 3 times in PBS and incubated in the corresponding secondary antibody (1:500; A32740, A32742, Invitrogen, Thermo Fisher Scientific. ab150169, Abcam) at 4°C overnight on a rotator. Slices were then washed three times with PBS, mounted with VECTASHIELD + DAPI and cover-slipped. All slices were imaged under a 2x objective on a fluorescence microscope (Keyence BZ-X810). Images were taken of 3 slices per mouse and fluorescence intensity was analyzed in ImageJ.

### Behavioral Studies

Spontaneous forepaw use was assessed by using the cylinder test at 3 and 6 weeks after unilateral injection. Mice were placed into a transparent plexiglass cylinder of 12 cm diameter and 30 cm height and video recorded on each side for a duration of 10 min. Rearing of the mice was analyzed for number of touches to the sides of the cylinder with either the paw ipsilateral to injection, the paw contralateral to injection or both forepaws simultaneously. Simultaneous forepaw use was excluded. Data was presented as percentage of the contralateral forepaw use by calculation with the equation: [(contralateral touches)/(ipsilateral touches + contralateral touches) × 100]. The calculated percentage declares the preference of the forepaw use as follows: 50% = symmetric use of both forepaws; <50% = preference of the ipsilateral forepaw; >50% preference of the contralateral forepaw.

### Blood collection and processing

At 42 days post-injection, blood was collected under isoflurane anesthesia (0.5–2% isoflurane in O_2_) via cardiac puncture with a 25-gauge needle attached to a heparinized syringe. Approximately 600-800 µL of blood was collected per mouse. Blood samples were immediately transferred to collection tubes containing either EDTA (for complete blood count, CBC) or serum separator tubes (for serum chemistry analysis). For CBC analysis, blood was analyzed within 4 hours of collection. The following parameters were measured: white blood cell count (WBC), red blood cell count (RBC), hemoglobin (Hb), hematocrit (Hct), mean corpuscular volume (MCV), platelets (PLT), and differential leukocyte counts (neutrophils, lymphocytes, monocytes, eosinophils, and basophils). For serum chemistry analysis, blood samples were allowed to clot at room temperature for 30 minutes. The samples were then centrifuged at 1,000 × g for 10 minutes at 4°C to separate the serum. The serum was transferred to a clean, labeled tube and stored at -80°C until analysis. The following parameters were measured: glucose, total protein, albumin, alkaline phosphatase (ALP), aspartate aminotransferase (AST), alanine aminotransferase (ALT), blood urea nitrogen (BUN), creatinine, total bilirubin, calcium, phosphorus, chloride, potassium, sodium.

### Statistical Analysis

For the cell culture experiments, the significance of the differences between no sgRNA and with sgRNA groups were analyzed statistically using student’s t test (Realstats Excel). Gene expression was measured via changes in luciferase concentration using a NanoLuc assay. A. A BCA Assay was also conducted to obtain the overall amino acid concentration and to normalize the data. The treated cells were then normalized using the control group of cells transfected/transduced with no sgRNA and no transcriptional repressor domain. Equations (1-2) demonstrate how the luciferase was normalized and quantified.

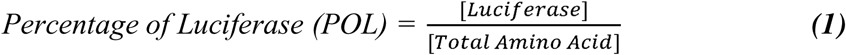

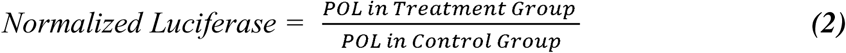

For the mouse experiments, all statistical analysis was performed using GraphPad Prism 10. To determine statistical significance, Shapiro-Wilk tests were first used to evaluate the assumption of normality of the data. For normally distributed data, one-way ANOVA, Two-way ANOVA or paired t-tests was used where appropriate, as described throughout the results.

## Data and materials availability

All data are available in the main text or the supplementary materials.

## Acknowledgments

National Institutes of Health/National Institute of Neurological Disorders and Stroke RF1 NS113548-01A1 (OC-F) Michael J. Fox Foundation for Parkinson’s Research MJFF-021362 (OC-F, BK)

## Author contributions

Conceptualization: OC-F, BK

Methodology: BOD, OC-F, BK

Investigation: BOD, JR, SU, DH

Funding acquisition: OC-F

Supervision: OC-F, BK

Writing – original draft: BO’D, O.C-F

Writing – review & editing: BOD, O.C-F, BK

