## Supplemental Table 1 for "*SNCA*-targeted epigenome therapy for Parkinson’s disease alleviates pathological and behavioral perturbations in a mouse model"

**Table. S1. Complete blood count and serum chemistry data**

| Parameter ($x10^3$) | Saline (mean±SEM) | | LV Repressor (mean±SEM) | | LV control (mean±SEM) | |
|---|---|---|---|---|---|---|
| | Male | Female | Male | Female | Male | Female |
| **Total WBCs** | 1.9 ± 0.2 | 0.85 ± 0.12 | 0.69 ± 0.15 | 0.69 ± 0.14 | 0.86 ± 0.17 | 0.63 ± 0.12 |
| **Neutrophils** | 0.2 ± 0.04 | 0.1 ± 0.02 | 0.06 ± 0.02 | 0.08 ± 0.02 | 0.12 ± 0.03 | 0.8 ± 0.01 |
| **Lymphocyte** | 1.6 ± 0.19 | 0.66 ± 0.12 | 0.59 ± 0.13 | 0.4 ± 0.12 | 0.7 ± 0.16 | 0.52 ± 0.1 |
| **Monocytes** | 0.06 ± 0.01 | 0.4 ± 0.01 | 0.02 ± 0.01 | 0.02 ± 0.01 | 0.03 ± 0.01 | 0.03 ± 0.01 |
| **Eosinophils** | 0.02 ± 0.005 | 0.01 ± 0.005 | 0.01 ± 0.005 | 0.002 ± 0.002 | 0.007 ± 0.003 | 0.007 ± 0.005 |
| **Basophils** | 0.002 ± 0.001 | 0.003 ± 0.002 | 0 | 0 | 0 | 0 |
| **RBCs** | 9.7 ± 0.06 | 8.6 ± 0.36 | 9.6 ± 0.14 | 8.7 ± 0.17 | 9.3 ± 0.24 | 9.4 ± 0.12 |
| **Platelets** | 1279 ± 149 | 1110 ± 154 | 1119 ± 241 | 970 ± 196 | 1221 ± 155 | 909 ± 72 |
| **Hematocrit** | 42.3 ± 0.35 | 42.05 ± 0.55 | 40.8 ± 0.55 | 40.7 ± 0.52 | 40.58 ± 0.95 | 40.58 ± 0.48 |
| **Hemoglobin** | 13.79 ± 0.1 | 13.75 ± 0.17 | 13.52 ± 0.15 | 13.57 ± 0.19 | 13.22 ± 0.24 | 13.4 ± 0.11 |
| **MCV** | 43.6 ± 0.12 | 43.7 ± 0.25 | 42.68 ± 1.08 | 44.8 ± 0.25 | 43.58 ± 0.38 | 43.38 ± 0.27 |
| **ALP** | 52.77 ± 1.19 | 71.67 ± 3.58 | 58.6 ± 8.63 | 75.83 ± 4.25 | 51.17 ± 2.74 | 73.3 ± 4.23 |
| **AST** | 65.89 ± 6.79 | 67.5 ± 10.99 | 68.6 ± 15.35 | 61.17 ± 6.54 | 51.5 ± 7.2 | 68.67 ± 7.86 |
| **ALT** | 7.22 ± 0.64 | 5.5 ± 0.85 | 7.8 ± 2.52 | 5 ± 0.58 | 4.33 ± 0.42 | 4.5 ± 0.43 |
| **BUN** | 26.78 ± 1.12 | 29.17 ± 0.19 | 29.8 ± 2.56 | 26.17 ± 0.6 | 25.17 ± 0.95 | 24.33 ± 0.67 |
| **Total protein** | 4.5 ± 0.04 | 4.33 ± 0.13 | 4.32 ± 0.89 | 4.08 ± 0.09 | 4.35 ± 0.06 | 4.03 ± 0.07 |
