## Supplemental Figure 1 for "*SNCA*-targeted epigenome therapy for Parkinson’s disease alleviates pathological and behavioral perturbations in a mouse model"

**Figure S1. Plasmid Sequences**

1. **pBK1838 pAAV-CMV-SNCAintron1-AlphaSyn[A53T]-WPRE-SV40pA (PD-INDUCER)**

cctgcaggcagctgcgcgctcgctcgctcactgaggccgcccgggcgtcgggcgacctttggtcgcccggcctcagtgagcgagcgag
cgcgcagagagggagtggccaactccatcactaggggttcctgcggccgcacgcgtATcgataagcttgggagttccgcgttacataact
tacggtaaatggcccgcctggctgaccgcccaacgacccccgcccattgacgtcaataatgacgtatgttcccatagtaacgccaataggg
actttccattgacgtcaatgggtggagtatttacggtaaactgcccacttggcagtacatcaagtgtatcatatgccaagtacgccccctattga
cgtcaatgacggtaaatggcccgcctggcattatgcccagtacatgaccttatgggactttcctacttggcagtacatctacgtattagtcatcg
ctattaccatggtgatgcggttttggcagtacatcaatgggcgtggatagcggtttgactcacggggatttccaagtctccaccccattgacgt
caatgggagtttgttttggcaccaaaatcaacgggactttccaaaatgtcgtaacaactccgccccattgacgcaaatgggcggtaggcgtgt
acggtgggaggtctatataagcagagctctctggctaactagagaacccactgcttactggcttatcgaaattaatacgactcactataggga
gacccaagctggCTAGCcgggcgtctttggaaatcctggagaacgccggatgggagacgaatggtcgtgggcaccgggagggggt
ggtgctgccatgaggacccgctgggccaggtctctgggaggtgagtacttgtccctttggggagcctaaggaaagagacttgacctggctt
tcgtcctgcttctgatattcccttctccacaagggctgagagattaggctgcttctccgggatccgcttttccccgggaaacgcgaggatgctc
catggagcgtgagcatccaacttttctctcacataaaatctgtctgcccgctctcttggtttttctctgtaaagtaagcaagctgcgtttggcaaat
aatgaaatggaagtgcaaggaggccaagtcaacaggtggtaacgggttaacaagtgctggcgcggggtccgctagggtggaggctgag
aacgccccctcgggtggctggcgcggggttggagacggcccgcgagtgtgagcggcgcctgctcagggtagatagctgagggcggCt
cgagctcaagcttcgatggatgtattcatgaaaggactttcaaaggccaaggagggagttgtggctgctgctgagaaaaccaaacagggtg
tggcagaagcagcaggaaagacaaaagagggtgttctctatgtaggctccaaaaccaaggagggagtggtgcatggtgtgacaacagtg
gctgagaagaccaaagagcaagtgacaaatgttggaggagcagtggtgacgggtgtgacagcagtagcccagaagacagtggaggga
gcagggagcattgcagcagccactggctttgtcaaaaaggaccagttgggcaagaatgaagaaggagccccacaggaaggaattctgga
agatatgcctgtggatcctgacaatgaggcttatgaaatgccttctgaggaagggtatcaagactacgaacctgaagcctaagaattctgcag
tcgagcgggatcaattccgccccccccctaacgttactggccgaagccgcttggaataaggccggtgtgcgtttgtctatatgttattttccac
catattgccgtcttttggcaatgtgagggcccggaaacctggccctgtcttcttgacgagcattcctaggggtctttcccctctcgccaaagga
atgcaaggtctgttgaatgtcgtgaaggaagcagttcctctggaagcttcttgaagacaaacaacgtctgtagcgaccctttgcaggcagcg
gaaccccccacctggcgacaggtgcctctgcggccaaaagccacgtgtataagatacacctgcaaaggcggcacaaccccagtgccac
gttgtgagttggatagttgtggaaagagtcaaatggctctcctcaagcgtattcaacaaggggctgaaggatgcccagaaggtacaagtaag
cGGCCGCTccggaatcaacctctggattacaaaatttgtgaaagattgactggtattcttaactatgttgctccttttacgctatgtggatac
gctgctttaatgcctttgtatcatgctattgcttcccgtatggctttcattttctcctccttgtataaatcctggttgctgtctctttatgaggagttgtgg
cccgttgtcaggcaacgtggcgtggtgtgcactgtgtttgctgacgcaacccccactggttggggcattgccaccacctgtcagctcctttcc
gggactttcgctttccccctccctattgccacggcggaactcatcgccgcctgccttgcccgctgctggacaggggctcggctgttgggcac
tgacaattccgtggtgttgtcggggaagctgacgtcctttccatggctgctcgcctgtgttgccacctggattctgcgcgggacgtccttctgct
acgtcccttcggccctcaatccagcggaccttccttcccgcggcctgctgccggctctgcggcctcttccgcgtcttcgccttcgccctcaga
cgagtcggatctccctttgggccgcctccccgcctgGATCCgtcgacccgggcggccgcttcgagcagacatgataagatacattgat
gagtttggacaaaccacaactagaatgcagtgaaaaaaatgctttatttgtgaaatttgtgatgctattgctttatttgtaaccattataagctgca
ataaacaagttaacaacaacaattgcattcattttatgtttcaggttcagggggagatgtgggaggttttttaaaGgtaaccacgtgcggaccg
agcggccgcaggaacccctagtgatggagttggccactccctctctgcgcgctcgctcgctcactgaggccgggcgaccaaaggtcgcc
cgacgcccgggctttgcccgggcggcctcagtgagcgagcgagcgcgcagctgcctgcaggggcgcctgatgcggtattttctccttacg
catctgtgcggtatttcacaccgcatacgtcaaagcaaccatagtacgcgccctgtagcggcgcattaagcgcggcgggtgtggtggttac
gcgcagcgtgaccgctacacttgccagcgccttagcgcccgctcctttcgctttcttcccttcctttctcgccacgttcgccggctttccccgtc

aagctctaaatcgggggctccctttagggttccgatttagtgctttacggcacctcgaccccaaaaaacttgatttgggtgatggttcacgtagt
gggccatcgccctgatagacggtttttcgccctttgacgttggagtccacgttctttaatagtggactcttgttccaaactggaacaacactcaa
ctctatctcgggctattcttttgatttataagggattttgccgatttcggtctattggttaaaaaatgagctgatttaacaaaaatttaacgcgaatttt
aacaaaatattaacgtttacaattttatggtgcactctcagtacaatctgctctgatgccgcatagttaagccagccccgacacccgccaacac
ccgctgacgcgccctgacgggcttgtctgctcccggcatccgcttacagacaagctgtgaccgtctccgggagctgcatgtgtcagaggttt
tcaccgtcatcaccgaaacgcgcgagacgaaagggcctcgtgatacgcctatttttataggttaatgtcatgataataatggtttcttagacgtc
aggtggcacttttcggggaaatgtgcgcggaacccctatttgtttatttttctaaatacattcaaatatgtatccgctcatgagacaataaccctga
taaatgcttcaataatattgaaaaaggaagagtatgagtattcaacatttccgtgtcgcccttattcccttttttgcggcattttgccttcctgtttttg
ctcacccagaaacgctggtgaaagtaaaagatgctgaagatcagttgggtgcacgagtgggttacatcgaactggatctcaacagcggtaa
gatccttgagagttttcgccccgaagaacgttttccaatgatgagcacttttaaagttctgctatgtggcgcggtattatcccgtattgacgccgg
gcaagagcaactcggtcgccgcatacactattctcagaatgacttggttgagtactcaccagtcacagaaaagcatcttacggatggcatga
cagtaagagaattatgcagtgctgccataaccatgagtgataacactgcggccaacttacttctgacaacgatcggaggaccgaaggagct
aaccgcttttttgcacaacatgggggatcatgtaactcgccttgatcgttgggaaccggagctgaatgaagccataccaaacgacgagcgtg
acaccacgatgcctgtagcaatggcaacaacgttgcgcaaactattaactggcgaactacttactctagcttcccggcaacaattaatagact
ggatggaggcggataaagttgcaggaccacttctgcgctcggcccttccggctggctggtttattgctgataaatctggagccggtgagcgt
gggtctcgcggtatcattgcagcactggggccagatggtaagccctcccgtatcgtagttatctacacgacggggagtcaggcaactatgg
atgaacgaaatagacagatcgctgagataggtgcctcactgattaagcattggtaactgtcagaccaagtttactcatatatactttagattgatt
taaaacttcatttttaatttaaaaggatctaggtgaagatcctttttgataatctcatgaccaaaatcccttaacgtgagttttcgttccactgagcgt
cagaccccgtagaaaagatcaaaggatcttcttgagatcctttttttctgcgcgtaatctgctgcttgcaaacaaaaaaaccaccgctaccagc
ggtggtttgtttgccggatcaagagctaccaactctttttccgaaggtaactggcttcagcagagcgcagataccaaatactgttcttctagtgt
agccgtagttaggccaccacttcaagaactctgtagcaccgcctacatacctcgctctgctaatcctgttaccagtggctgctgccagtggcg
ataagtcgtgtcttaccgggttggactcaagacgatagttaccggataaggcgcagcggtcgggctgaacggggggttcgtgcacacagc
ccagcttggagcgaacgacctacaccgaactgagatacctacagcgtgagctatgagaaagcgccacgcttcccgaagggagaaaggc
ggacaggtatccggtaagcggcagggtcggaacaggagagcgcacgagggagcttccagggggaaacgcctggtatctttatagtcctg
tcgggtttcgccacctctgacttgagcgtcgatttttgtgatgctcgtcaggggggcggagcctatggaaaaacgccagcaacgcggccttttt
tacggttcctggccttttgctggccttttgctcacatgt

2. **pBK1132- Lenti-EFSNC-dSaCas9-U6-SAgRNA scaffold (INACTIVE VECTOR, no gRNA, no repressor)**

gcacttttcggggaaatgtgcgcggaacccctatttgtttatttttctaaatacattcaaatatgtatccgctcatgagacaataaccctgataaat
gcttcaataatattgaaaaaggaagagtatgagtattcaacatttccgtgtcgcccttattcccttttttgcggcattttgccttcctgtttttgctcac
ccagaaacgctggtgaaagtaaaagatgctgaagatcagttgggtgcacgagtgggttacatcgaactggatctcaacagcggtaagatcc
ttgagagttttcgccccgaagaacgttttccaatgatgagcacttttaaagttctgctatgtggcgcggtattatcccgtattgacgccgggcaa
gagcaactcggtcgccgcatacactattctcagaatgacttggttgagtactcaccagtcacagaaaagcatcttacggatggcatgacagta
agagaattatgcagtgctgccataaccatgagtgataacactgcggccaacttacttctgacaacgatcggaggaccgaaggagctaaccg
cttttttgcacaacatgggggatcatgtaactcgccttgatcgttgggaaccggagctgaatgaagccataccaaacgacgagcgtgacacc
acgatgcctgtagcaatggcaacaacgttgcgcaaactattaactggcgaactacttactctagcttcccggcaacaattaatagactggatg
gaggcggataaagttgcaggaccacttctgcgctcggcccttccggctggctggtttattgctgataaatctggagccggtgagcgtgggtc
tcgcggtatcattgcagcactggggccagatggtaagccctcccgtatcgtagttatctacacgacggggagtcaggcaactatggatgaa

cgaaatagacagatcgctgagataggtgcctcactgattaagcattggtaactgtcagaccaagtttactcatatatactttagattgatttaaaa
cttcatttttaatttaaaaggatctaggtgaagatcctttttgataatctcatgaccaaaatcccttaacgtgagttttcgttccactgagcgtcaga
ccccgtagaaaagatcaaaggatcttcttgagatcctttttttctgcgcgtaatctgctgcttgcaaacaaaaaaaccaccgctaccagcggtg
gtttgtttgccggatcaagagctaccaactctttttccgaaggtaactggcttcagcagagcgcagataccaaatactgttcttctagtgtagcc
gtagttaggccaccacttcaagaactctgtagcaccgcctacatacctcgctctgctaatcctgttaccagtggctgctgccagtggcgataa
gtcgtgtcttaccgggttggactcaagacgatagttaccggataaggcgcagcggtcgggctgaacggggggttcgtgcacacagcccag
cttggagcgaacgacctacaccgaactgagatacctacagcgtgagctatgagaaagcgccacgcttcccgaagggagaaaggcggac
aggtatccggtaagcggcagggtcggaacaggagagcgcacgagggagcttccagggggaaacgcctggtatctttatagtcctgtcgg
gtttcgccacctctgacttgagcgtcgatttttgtgatgctcgtcaggggggcggagcctatggaaaaacgccagcaacgcggcctttttacg
gttcctggccttttgctggccttttgctcacatgttctttcctgcgttatcccctgattctgtggataaccgtattaccgcctttgagtgagctgatac
cgctcgccgcagccgaacgaccgagcgcagcgagtcagtgagcgaggaagcggaagagcgcccaatacgcaaaccgcctctccccg
cgcgttggccgattcattaatgcagctggcacgacaggtttcccgactggaaagcgggcagtgagcgcaacgcaattaatgtgagttagct
cactcattaggcaccccaggctttacactttatgcttccggctcgtatgttgtgtggaattgtgagcggataacaatttcacacaggaaacagct
atgaccatgattacgccaagcgcgcaattaaccctcactaaagggaacaaaagctggagctgcaagcttaatgtagtcttatgcaatactctt
gtagtcttgcaacatggtaacgatgagttagcaacatgccttacaaggagagaaaaagcaccgtgcatgccgattggtggaagtaaggtgg
tacgatcgtgccttattaggaaggcaacagacgggtctgacatggattggacgaaccactgaattgccgcattgcagagatattgtatttaagt
gcctagctcgatacataaacgggtctctctggttagaccagatctgagcctgggagctctctggctaactagggaacccactgcttaagcctc
aataaagcttgccttgagtgcttcaagtagtgtgtgcccgtctgttgtgtgactctggtaactagagatccctcagacccttttagtcagtgtgga
aaatctctagcagtggcgcccgaacagggacttgaaagcgaaagggaaaccagaggagctctctcgacgcaggactcggcttgctgaag
cgcgcacggcaagaggcgaggggcggcgactggtgagtacgccaaaaattttgactagcggaggctagaaggagagagatgggtgcg
agagcgtcagtattaagcgggggagaattagatcgcgatgggaaaaaattcggttaaggccagggggaaagaaaaaatataaattaaaac
atatagtatgggcaagcagggagctagaacgattcgcagttaatcctggcctgttagaaacatcagaaggctgtagacaaatactgggaca
gctacaaccatcccttcagacaggatcagaagaacttagatcattatataatacagtagcaaccctctattgtgtgcatcaaaggatagagata
aaagacaccaaggaagctttagacaagatagaggaagagcaaaacaaaagtaagaccaccgcacagcaagcggccgctgatcttcaga
cctggaggaggagatatgagggacaattggagaagtgaattatataaatataaagtagtaaaaattgaaccattaggagtagcacccaccaa
ggcaaagagaagagtggtgcagagagaaaaaagagcagtgggaataggagctttgttccttgggttcttgggagcagcaggaagcactat
gggcgcagcgtcaatgacgctgacggtacaggccagacaattattgtctggtatagtgcagcagcagaacaatttgctgagggctattgag
gcgcaacagcatctgttgcaactcacagtctggggcatcaagcagctccaggcaagaatcctggctgtggaaagatacctaaaggatcaa
cagctcctggggatttggggttgctctggaaaactcatttgcaccactgctgtgccttggaatgctagttggagtaataaatctctggaacagat
ttggaatcacacgacctggatggagtgggacagagaaattaacaattacacaagcttaatacactccttaattgaagaatcgcaaaaccagc
aagaaaagaatgaacaagaattattggaattagataaatgggcaagtttgtggaattggtttaacataacaaattggctgtggtatataaaattat
tcataatgatagtaggaggcttggtaggtttaagaatagtttttgctgtactttctatagtgaatagagttaggcagggatattcaccattatcgttt
cagacccacctcccaaccccgaggggacccgacaggcccgaaggaatagaagaagaaggtggagagagagacagagacagatccat
tcgattagtgaacggatctcgacggtatcggttaacttttaaaagaaaaggggggattggggggtacagtgcaggggaaagaatagtagac
ataatagcaacagacatacaaactaaagaattacaaaaacaaattacaaaattcaaaattttatcgGcttgaaaggagtgggaattggctccg
gtgcccgtcagtgggcagagcgcacatcgcccacagtccccgagaagttggggggaggggtcggcaattgatccggtgcctagagaag
gtggcgcggggtaaactgggaaagtgatgtcgtgtactggctccgcctttttcccgagggtgggggagaaccgtatataagtgcagtagtc
gccgtgaacgttctttttcgcaacgggtttgccgccagaacacaggaccggtgccaccatgGACTACAAAGACCATGACG
GTGATTATAAAGATCATGACATaGATTACAAGGATGACGATGACAAGggtggatcgccaaagaaa
aagaggaaagtcgccccaaagaagaagcggaaggtcggtatccacggagtcccagcagccaagcggaactacatcctgggcctggcc

atcggcatcaccagcgtgggctacggcatcatcgactacgagacacgggacgtgatcgatgccggcgtgcggctgttcaaagaggccaa cgtggaaaacaacgagggcaggcggagcaagagaggcgccagaaggctgaagcggcggaggcggcatagaatccagagagtgaag aagctgctgttcgactacaacctgctgaccgaccacagcgagctgagcggcatcaacccctacgaggccagagtgaagggcctgagcca gaagctgagcgaggaagagttctctgccgccctgctgcacctggccaagagaagaggcgtgcacaacgtgaacgaggtggaagagga caccggcaacgagctgtccaccaaagagcagatcagccggaacagcaaggccctggaagagaaatacgtggccgaactgcagctgga acggctgaagaaagacggcgaagtgcggggcagcatcaacagattcaagaccagcgactacgtgaaagaagccaaacagctgctgaa ggtgcagaaggcctaccaccagctggaccagagcttcatcgacacctacatcgacctgctggaaacccggcggacctactatgagggac ctggcgagggcagccccttcggctggaaggacatcaaagaatggtacgagatgctgatgggccactgcacctacttccccgaggaactg cggagcgtgaagtacgcctacaacgccgacctgtacaacgccctgaacgacctgaacaatctcgtgatcaccagggacgagaacgagaa gctggaatattacgagaagttccagatcatcgagaacgtgttcaagcagaagaagaagcccaccctgaagcagatcgccaaagaaatcct cgtgaacgaagaggatattaagggctacagagtgaccagcaccggcaagcccgagttcaccaacctgaaggtgtaccacgacatcaagg acattaccgcccggaaagagattattgagaacgccgagctgctggatcagattgccaagatcctgaccatctaccagagcagcgaggaca tccaggaagaactgaccaatctgaactccgagctgacccaggaagagatcgagcagatctctaatctgaagggctataccggcacccaca acctgagcctgaaggccatcaacctgatcctggacgagctgtggcacaccaacgacaaccagatcgctatcttcaaccggctgaagctggt gcccaagaaggtggacctgtcccagcagaaagagatccccaccaccctggtggacgacttcatcctgagccccgtcgtgaagagaagctt catccagagcatcaaagtgatcaacgccatcatcaagaagtacggcctgcccaacgacatcattatcgagctggcccgcgagaagaactc caaggacgcccagaaaatgatcaacgagatgcagaagcggaaccggcagaccaacgagcggatcgaggaaatcatccggaccaccg gcaaagagaacgccaagtacctgatcgagaagatcaagctgcacgacatgcaggaaggcaagtgcctgtacagcctggaagccatccct ctggaagatctgctgaacaaccccttcaactatgaggtggaccacatcatccccagaagcgtgtccttcgacaacagcttcaacaacaaggt gctcgtgaagcaggaagaagccagcaagaagggcaaccggaccccattccagtacctgagcagcagcgacagcaagatcagctacga aaccttcaagaagcacatcctgaatctggccaagggcaagggcagaatcagcaagaccaagaaagagtatctgctggaagaacgggac atcaacaggttctccgtgcagaaagacttcatcaaccggaacctggtggataccagatacgccaccagaggcctgatgaacctgctgcgg agctacttcagagtgaacaacctggacgtgaaagtgaagtccatcaatggcggcttcaccagctttctgcggcggaagtggaagtttaagaa agagcggaacaaggggtacaagcaccacgccgaggacgccctgatcattgccaacgccgatttcatcttcaaagagtggaagaaactgg acaaggccaaaaaagtgatggaaaaccagatgttcgaggaaaagcaggccgagagcatgcccgagatcgaaaccgagcaggagtaca aagagatcttcatcacccccccaccagatcaagcacattaaggacttcaaggactacaagtacagccaccgggtggacaagaagcctaata gagagctgattaacgacaccctgtactccacccggaaggacgacaagggcaacaccctgatcgtgaacaatctgaacggcctgtacgac aaggacaatgacaagctgaaaaagctgatcaacaagagccccgaaaagctgctgatgtaccaccacgacccccagacctaccagaaact gaagctgattatggaacagtacggcgacgagaagaatcccctgtacaagtactacgaggaaaccgggaactacctgaccaagtactccaa aaaggacaacggccccgtgatcaagaagattaagtattacggcaacaaactgaacgcccatctggacatcaccgacgactaccccaacag cagaaacaaggtcgtgaagctgtccctgaagccctacagattcgacgtgtacctggacaatggcgtgtacaagttcgtgaccgtgaagaat ctggatgtgatcaaaaaagaaaactactacgaagtgaatagcaagtgctatgaggaagctaagaagctgaagaagatcagcaaccaggcc gagtttatcgcctccttctacaacaacgatctgatcaagatcaacggcgagctgtatagagtgatcggcgtgaacaacgacctgctgaaccg gatcgaagtgaacatgatcgacatcacctaccgcgagtacctggaaaacatgaacgacaagaggcccccccaggatcattaagacaatcgc ctccaagacccagagcattaagaagtacagcacagacattctgggcaacctgtatgaagtgaaatctaagaagcaccctcagatcatcaaa aagggcaaaaggccggcggccacgaaaaaggccggccaggcaaaaaagaaaaaggGATCCTGAAATAAAAGATCT TTGTTTTCATTAGATCTGTGTGTTGGTTTTTTGTGTGCggCCggtaccgagggcctatttcccatgattcc ttcatatttgcatatacgatacaaggctgttagagagataattggaattaatttgactgtaaacacaaagatattagtacaaaatacgtgacgtag aaagtaataatttcttgggtagtttgcagttttaaaattatgttttaaaatggactatcatatgcttaccgtaacttgaaagtatttcgatttcttggcttt atatatcttgtggaaaggacgaaacaccggagaccacggcaggtctcagttttagtactctggaaacagaatctactaaaacaaggcaaaat

gccgtgtttatctcgtcaacttgttggcgagattAGCaattcgagctcggtacctttaagaccaatgacttacaaggcagctgtagatcttag
ccactttttaaaagaaaaggggggactggaagggctaattcactcccaacgaagacaagatctgctttttgcttgtactgggtctctctggtta
gaccagatctgagcctgggagctctctggctaactagggaacccactgcttaagcctcaataaagcttgccttgagtgcttcaagtagtgtgt
gcccgtctgttgtgtgactctggtaactagagatccctcagacccttttagtcagtgtggaaaatctctagcagtagtagttcatgtcatcttatta
ttcagtatttataacttgcaaagaaatgaatatcagagagtgagaggaacttgtttattgcagcttataatggttacaaataaagcaatagcatca
caaatttcacaaataaagcatttttttcactgcattctagttgtggtttgtccaaactcatcaatgtatcttatcatgtctggctctagctatcccgccc
ctaactccgcccatcccgcccctaactccgcccagttccgcccattctccgccccatggctgactaattttttttatttatgcagaggccgaggc
cgcctcggcctctgagctattccagaagtagtgaggaggcttttttggaggcctagggacgtacccaattcgccctatagtgagtcgtattac
gcgcgctcactggccgtcgttttacaacgtcgtgactgggaaaaccctggcgttacccaacttaatcgccttgcagcacatccccctttcgcc
agctggcgtaatagcgaagaggcccgcaccgatcgcccttcccaacagttgcgcagcctgaatggcgaatgggacgcgccctgtagcgg
cgcattaagcgcggcgggtgtggtggttacgcgcagcgtgaccgctacacttgccagcgccctagcgcccgctcctttcgctttcttcccttc
ctttctcgccacgttcgccggctttccccgtcaagctctaaatcgggggctccctttagggttccgatttagtgctttacggcacctcgacccca
aaaaacttgattagggtgatggttcacgtagtgggccatcgccctgatagacggtttttcgccctttgacgttggagtccacgttctttaatagtg
gactcttgttccaaactggaacaacactcaaccctatctcggtctattcttttgatttataagggattttgccgatttcggcctattggttaaaaaat
gagctgatttaacaaaaatttaacgcgaattttaacaaaatattaacgcttacaatttaggtg

### 3. pBK1579- pLV-U6-[empty]-EFSNC-dSaCas9-DNMT3a-P2A-Puro-WPRE (pCL17) (no gRNA vector)

ctagttattaatagtaatcaattacggggtcattagttcatagcccatatatggagttccgcgttacataacttacggtaaatggcccgcctggct
gaccgcccaacgacccccgcccattgacgtcaataatgacgtatgttcccatagtaacgccaatagggactttccattgacgtcaatgggtg
gagtatttacggtaaactgcccacttggcagtacatcaagtgtatcatatgccaagtacgccccctattgacgtcaatgacggtaaatggccc
gcctggcattatgcccagtacatgaccttatgggactttcctacttggcagtacatctacgtattagtcatcgctattaccatggtgatgcggtttt
ggcagtacatcaatgggcgtggatagcggtttgactcacggggatttccaagtctccaccccattgacgtcaatgggagtttgttttggcacc
aaaatcaacgggactttccaaaatgtcgtaacaactccgccccattgacgcaaatgggcggtaggcgtgtacggtgggaggtctatataag
cagcgcgttttgcctgtactgggtctctctggttagaccagatctgagcctgggagctctctggctaactagggaacccactgcttaagcctca
ataaagcttgccttgagtgcttcaagtagtgtgtgcccgtctgttgtgtgactctggtaactagagatccctcagacccttttagtcagtgtggaa
aatctctagcagtggcgcccgaacagggacttgaaagcgaaagggaaaccagaggagctctctcgacgcaggactcggcttgctgaagc
gcgcacggcaagaggcgaggggcggcgactggtgagtacgccaaaaattttgactagcggaggctagaaggagagagatgggtgcga
gagcgtcagtattaagcgggggagaattagatcgcgatgggaaaaaattcggttaaggccagggggaaagaaaaaatataaattaaaaca
tatagtatgggcaagcagggagctagaacgattcgcagttaatcctggcctgttagaaacatcagaaggctgtagacaaatactgggacag
ctacaaccatcccttcagacaggatcagaagaacttagatcattatataatacagtagcaaccctctattgtgtgcatcaaaggatagagataa
aagacaccaaggaagctttagacaagatagaggaagagcaaaacaaaagtaagaccaccgcacagcaagcggccgctgatcttcagac
ctggaggaggagatatgagggacaattggagaagtgaattatataaatataaagtagtaaaaattgaaccattaggagtagcacccaccaa
ggcaaagagaagagtggtgcagagagaaaaaagagcagtgggaataggagctttgttccttgggttcttgggagcagcaggaagcactat
gggcgcagcgtcaatgacgctgacggtacaggccagacaattattgtctggtatagtgcagcagcagaacaatttgctgagggctattgag
gcgcaacagcatctgttgcaactcacagtctggggcatcaagcagctccaggcaagaatcctggctgtggaaagatacctaaaggatcaa
cagctcctggggatttggggttgctctggaaaactcatttgcaccactgctgtgccttggaatgctagttggagtaataaatctctggaacagat
ttggaatcacacgacctggatggagtgggacagagaaattaacaattacacaagcttaatacactccttaattgaagaatcgcaaaaccagc
aagaaaagaatgaacaagaattattggaattagataaatgggcaagtttgtggaattggtttaacataacaaattggctgtggtatataaaattat

tcataatgatagtaggaggcttggtaggtttaagaatagtttttgctgtactttctatagtgaatagagttaggcagggatattcaccattatcgttt
cagacccacctcccaaccccgaggggacccgacaggcccgaaggaatagaagaagaaggtggagagagagacagagacagatccat
tcgattagtgaacggatcggcactgcgtgcgccaattctgcagacaaatggcagtattcatccacaattttaaaagaaaaggggggattggg
gggtacagtgcaggggaaagaatagtagacataatagcaacagacatacaaactaaagaattacaaaaacaaattacaaaaattcaaaattt
tcgggtttattacagggacagcagagatccagtttggTTAATTAATGGGCGGGACGTTAACGGGGCGGAAC
GGTACCgagggcctatttcccatgattccttcatatttgcatatacgatacaaggctgttagagagataattagaattaatttgactgtaaac
acaaagatattagtacaaaatacgtgacgtagaaagtaataatttcttgggtagtttgcagttttaaaattatgttttaaaatggactatcatatgctt
accgtaacttgaaagtatttcgatttcttggctttatatatcttGTGGAAAGGACGAAAcaccggagacgtgtacacgtctctgT
TTtagtactctggaaacagaatctactaaaacaaggcaaaatgccgtgtttatctcgtcaacttgttggcgagaTTTTTTgaattcgcta
gctaggtcttgaaaggagtgggaattggctccggtgcccgtcagtgggcagagcgcacatcgcccacagtccccgagaagttggggga
ggggtcggcaattgatccggtgcctagagaaggtggcgcggggtaaactgggaaagtgatgtcgtgtactggctccgcctttttcccgagg
gtgggggagaaccgtatataagtgcagtagtcgccgtgaacgttctttttcgcaacgggtttgccgccagaacacaggaccggtgccacca
tgGACTACAAAGACCATGACGGTGATTATAAAGATCATGACATaGATTACAAGGATGAC
GATGACAAGggtggatcgccaaagaaaaagaggaaagtcgccccaaagaagaagcggaaggtcggtatccacggagtcccag
cagccaagcggaactacatcctgggcctggccatcggcatcaccagcgtgggctacggcatcatcgactacgagacacgggacgtgatc
gatgccggcgtgcggctgttcaaagaggccaacgtggaaaacaacgagggcaggcggagcaagagaggcgccagaaggctgaagcg
gcggaggcggcatagaatccagagagtgaagaagctgctgttcgactacaacctgctgaccgaccacagcgagctgagcggcatcaac
ccctacgaggccagagtgaagggcctgagccagaagctgagcgaggaagagttctctgccgccctgctgcacctggccaagagaagag
gcgtgcacaacgtgaacgaggtggaagaggacaccggcaacgagctgtccaccaaagagcagatcagccggaacagcaaggccctg
gaagagaaatacgtggccgaactgcagctggaacggctgaagaaagacggcgaagtgcggggcagcatcaacagattcaagaccagc
gactacgtgaaagaagccaaacagctgctgaaggtgcagaaggcctaccaccagctggaccagagcttcatcgacacctacatcgacct
gctggaaacccggcggacctactatgagggacctggcgagggcagccccttcggctggaaggacatcaaagaatggtacgagatgctg
atgggccactgcacctacttccccgaggaactgcggagcgtgaagtacgcctacaacgccgacctgtacaacgccctgaacgacctgaa
caatctcgtgatcaccagggacgagaacgagaagctggaatattacgagaagttccagatcatcgagaacgtgttcaagcagaagaagaa
gcccaccctgaagcagatcgccaaagaaatcctcgtgaacgaagaggatattaagggctacagagtgaccagcaccggcaagcccgag
ttcaccaacctgaaggtgtaccacgacatcaaggacattaccgcccggaaagagattattgagaacgccgagctgctggatcagattgcca
agatcctgaccatctaccagagcagcgaggacatccaggaagaactgaccaatctgaactccgagctgacccaggaagagatcgagcag
atctctaatctgaagggctataccggcacccacaacctgagcctgaaggccatcaacctgatcctggacgagctgtggcacaccaacgac
aaccagatcgctatcttcaaccggctgaagctggtgcccaagaaggtggacctgtcccagcagaaagagatccccaccaccctggtggac
gacttcatcctgagccccgtcgtgaagagaagcttcatccagagcatcaaagtgatcaacgccatcatcaagaagtacggcctgcccaacg
acatcattatcgagctggcccgcgagaagaactccaaggacgcccagaaaatgatcaacgagatgcagaagcggaaccggcagaccaa
cgagcggatcgaggaaatcatccggaccaccggcaaagagaacgccaagtacctgatcgagaagatcaagctgcacgacatgcaggaa
ggcaagtgcctgtacagcctggaagccatccctctggaagatctgctgaacaaccccttcaactatgaggtggaccacatcatccccagaa
gcgtgtccttcgacaacagcttcaacaacaaggtgctcgtgaagcaggaagaagccagcaagaagggcaaccggaccccattccagtac
ctgagcagcagcgacagcaagatcagctacgaaaccttcaagaagcacatcctgaatctggccaagggcaagggcagaatcagcaaga
ccaagaaagagtatctgctggaagaacgggacatcaacaggttctccgtgcagaaagacttcatcaaccggaacctggtggataccagat
acgccaccagaggcctgatgaacctgctgcggagctacttcagagtgaacaacctggacgtgaaagtgaagtccatcaatggcggcttca
ccagctttctgcggcggaagtggaagtttaagaaagagcggaacaaggggtacaagcaccacgccgaggacgccctgatcattgccaac
gccgatttcatcttcaaagagtggaagaaactggacaaggccaaaaaagtgatggaaaaccagatgttcgaggaaaagcaggccgagag
catgcccgagatcgaaaccgagcaggagtacaaagagatcttcatcaccccccaccagatcaagcacattaaggacttcaaggactacaa

gtacagccaccgggtggacaagaagcctaatagagagctgattaacgacaccctgtactccacccggaaggacgacaagggcaacacc
ctgatcgtgaacaatctgaacggcctgtacgacaaggacaatgacaagctgaaaaagctgatcaacaagagccccgaaaagctgctgatg
taccaccacgacccccagacctaccagaaactgaagctgattatggaacagtacggcgacgagaagaatcccctgtacaagtactacgag
gaaaccgggaactacctgaccaagtactccaaaaaggacaacggccccgtgatcaagaagattaagtattacggcaacaaactgaacgc
ccatctggacatcaccgacgactaccccaacagcagaaacaaggtcgtgaagctgtccctgaagccctacagattcgacgtgtacctgga
caatggcgtgtacaagttcgtgaccgtgaagaatctggatgtgatcaaaaaagaaaactactacgaagtgaatagcaagtgctatgaggaa
gctaagaagctgaagaagatcagcaaccaggccgagtttatcgcctccttctacaacaacgatctgatcaagatcaacggcgagctgtata
gagtgatcggcgtgaacaacgacctgctgaaccggatcgaagtgaacatgatcgacatcacctaccgcgagtacctggaaaacatgaacg
acaagaggcccccccaggatcattaagacaatcgcctccaagacccagagcattaagaagtacagcacagacattctgggcaacctgtatg
aagtgaaatctaagaagcaccctcagatcatcaaaaagggcaaaaggccggcggccacgaaaaaggcGcTCGAGGGCGGA
GGCGGGAGCGGATCCCCCTCCCGGCTCCAGATGttcttcgctaataaccacgaccaggaatttgaccctccaa
aggtttacccacctgtcccagctgagaagaggaagcccatccgggtgctgtctctctttgatggaatcgctacagggctcctggtgctgaag
gacttgggcattcaggtggaccgctacattgcctcggaggtgtgtgaggactccatcacggtgggcatggtgcggcaccaggggaagatc
atgtacgtcggggacgtccgcagcgtcacacagaagcatatccaggagtggggcccattcgatctggtgattgggggcagtccctgcaat
gacctctccatcgtcaaccctgctcgcaagggcctctacgagggcactggccggctcttctttgagttctaccgcctcctgcatgatgcgcgg
cccaaggagggagatgatcgcccottcttctggctctttgagaatgtggtggccatgggcgttagtgacaagagggacatctcgcgatttctc
gagtccaaccctgtgatgattgatgccaaagaagtgtcagctgcacacagggcccgctacttctggggtaaccttcccggtatgaacaggc
cgttggcatccactgtgaatgataagctggagctgcaggagtgtctggagcatggcaggatagccaagttcagcaaagtgaggaccattac
tacgaggtcaaactccataaagcagggcaaaGACCAGCATTTTCCTGTGTTCATGAATGAGAAAGAGga
catcttatggtgcactgaaatggaaagggtatttggtttcccagtccactatactgacgtgtccaacatgagccgcttggcgaggcagagact
gctgggccggtcatggagcgtgccagtcatccgccacctcttcgctcCGCTGAAGGAGTATTTTGCGTGTGTGtcc
ggccggcccggatccGGCGCAACAAACTTCTCTCTGCTGAAACAAGCCGGAGATGTCGAAG
AGAATCCTGGACCGACCGAGTACAAGCCCACGGTGCGCCTCGCCACCCGCGACGAC
GTCCCCAGGGCCGTACGCACCCTCGCCGCCGCGTTCGCCGACTACCCCGCCACGCGC
CACACCGTCGATCCGGACCGCCACATCGAGCGGGTCACCGAGCTGCAAGAACTCTTC
CTCACGCGCGTCGGGCTCGACATCGGCAAGGTGTGGGTCGCGGACGACGGCGCCGC
GGTGGCGGTCTGGACCACGCCGGAGAGCGTCGAAGCGGGGGCGGTGTTCGCCGAGA
TCGGCCCGCGCATGGCCGAGTTGAGCGGTTCCCGGCTGGCCGCGCAGCAACAGATG
GAAGGCCTCCTGGCGCCGCACCGGCCCAAGGAGCCCGCGTGGTTCCTGGCCACCGT
CGGAGTCTCGCCCGACCACCAGGGCAAGGGTCTGGGCAGCGCCGTCGTGCTCCCCG
GAGTGGAGGCGGCCGAGCGCGCCGGGGTGCCCGCCTTCCTGGAGACCTCCGCGCCC
CGCAACCTCCCCTTCTACGAGCGGCTCGGCTTCACCGTCACCGCCGACGTCGAGGTG
CCCGAAGGACCGCGCACCTGGTGCATGACCCGCAAGCCCGGTGCCTGAACGCGTTA
AGTCGACAATCAACCTCTGGATTACAAAATTTGTGAAAGATTGACTGGTATTCTTAAC
TATGTTGCTCCTTTTACGCTATGTGGATACGCTGCTTTAATGCCTTTGTATCATGCTATT
GCTTCCCGTATGGCTTTCATTTTCTCCTCCTTGTATAAATCCTGGTTGCTGTCTCTTTAT
GAGGAGTTGTGGCCCGTTGTCAGGCAACGTGGCGTGGTGTGCACTGTGTTTGCTGAC
GCAACCCCCACTGGTTGGGGCATTGCCACCACCTGTCAGCTCCTTTCCGGGACTTTC
GCTTTCCCCCTCCCTATTGCCACGGCGGAACTCATCGCCGCCTGCCTTGCCCGCTGCT
GGACAGGGGCTCGGCTGTTGGGCACTGACAATTCCGTGGTGTTGTCGGGGAAATCAT

CGTCCTTTCCTTGGCTGCTCGCCTGTGTTGCCACCTGGATTCTGCGCGGGACGTCCTT
CTGCTACGTCCCTTCGGCCCTCAATCCAGCGGACCTTCCTTCCCGCGGCCTGCTGCCG
GCTCTGCGGCCTCTTCCGCGTCTTCGCCTTCGCCCTCAGACGAGTCGGATCTCCCTTT
GGGCCGCCTCCCCGCGTCGACTTTAAGACCAATGACTTACAAGGCAGCTGTAGATCTT
AGCCACTTTTTAAAAGAAAAGGGGGGACTGGAAGGGCTAATTCACTCCCAACGAAG
ACAAGATCTGCTTTTTGCTTGTACTGGGTCTCTCTGGTTAGACCAGATCTGAGCCTGG
GAGCTCTCTGGCTAACTAGGGAACCCACTGCTTAAGCCTCAATAAAGCTTGCCTTGA
GTGCTTCAAGTAGTGTGTGCCCGTCTGTTGTGTGACTCTGGTAACTAGAGATCCCTCA
GACCCTTTTAGTCAGTGTGGAAAATCTCTAGCAGggcccgtttaaacccgctgatcagcctcgactgtgcctt
ctagttgccagccatctgttgtttgcccctcccccgtgccttccttgaccctggaaggtgccactcccactgtcctttcctaataaaatgaggaa
attgcatcgcattgtctgagtaggtgtcattctattctggggggtggggtggggcaggacagcaagggggaggattgggaagacaatagca
ggcatgctggggatgcggtgggctctatggcttctgaggcggaaagaaccagctggggctctagggggtatccccacgcgccctgtagc
ggcgcattaagcgcggcgggtgtggtggttacgcgcagcgtgaccgctacacttgccagcgccctagcgcccgctcctttcgctttcttccc
ttcctttctcgccacgttcgccggctttccccgtcaagctctaaatcgggggctccctttagggttccgatttagtgctttacggcacctcgaccc
caaaaaacttgattagggtgatggttcacgtagtgggccatcgccctgatagacggtttttcgccctttgacgttggagtccacgttctttaata
gtggactcttgttccaaactggaacaacactcaaccctatctcggtctattcttttgatttataatggttacaaataaagcaatagcatcacaaattt
cacaaataaagcatttttttcactgcattctagttgtggtttgtccaaactcatcaatgtatcttatcatgtctgtataccgtcgacctctagctagag
cttggcgtaatcatggtcatagctgtttcctgtgtgaaattgttatccgctcacaattccacacaacatacgagccggaagcataaagtgtaaag
cctggggtgcctaatgagtgagctaactcacattaattgcgttgcgctcactgcccgctttccagtcgggaaacctgtcgtgccagctgcatta
atgaatcggccaacgcgcggggagaggcggtttgcgtattgggcgctcttccgcttcctcgctcactgactcgctgcgctcggtcgttcggc
tgcggcgagcggtatcagctcactcaaaggcggtaatacggttatccacagaatcaggggataacgcaggaaagaacatgtgagcaaaa
ggccagcaaaaggccaggaaccgtaaaaaggccgcgttgctggcgtttttccataggctccgcccccctgacgagcatcacaaaaatcga
cgctcaagtcagaggtggcgaaacccgacaggactataaagataccaggcgtttccccctggaagctccctcgtgcgctctcctgttccga
ccctgccgcttaccggatacctgtccgcctttctcccttcgggaagcgtggcgctttctcatagctcacgctgtaggtatctcagttcggtgtag
gtcgttcgctccaagctgggctgtgtgcacgaaccccccgttcagcccgaccgctgcgccttatccggtaactatcgtcttgagtccaaccc
ggtaagacacgacttatcgccactggcagcagccactggtaacaggattagcagagcgaggtatgtaggcggtgctacagagttcttgaa
gtggtggcctaactacggctacactagaagaacagtatttggtatctgcgctctgctgaagccagttaccttcggaaaaagagttggtagctct
tgatccggcaaacaaaccaccgctggtagcggtggtttttttgtttgcaagcagcagattacgcgcagaaaaaaaggatctcaagaagatcc
tttgatcttttctacggggtctgacgctcagtggaacgaaaactcacgttaagggattttggtcatgagattatcaaaaaggatcttcacctagat
ccttttaaattaaaaatgaagttttaaatcaatctaaagtatatatgagtaaacttggtctgacagttaccaatgcttaatcagtgaggcacctatct
cagcgatctgtctatttcgttcatccatagttgcctgactccccgtcgtgtagataactacgatacgggagggcttaccatctggccccagtgct
gcaatgataccgcgagacccacgctcaccggctccagatttatcagcaataaaccagccagccggaagggccgagcgcagaagtggtc
ctgcaactttatccgcctccatccagtctattaattgttgccgggaagctagagtaagtagttcgccagttaatagtttgcgcaacgttgttgcca
ttgctacaggcatcgtggtgtcacgctcgtcgtttggtatggcttcattcagctccggttcccaacgatcaaggcgagttacatgatcccccat
gttgtgcaaaaaagcggttagctccttcggtcctccgatcgttgtcagaagtaagttggccgcagtgttatcactcatggttatggcagcactg
cataattctcttactgtcatgccatccgtaagatgcttttctgtgactggtgagtactcaaccaagtcattctgagaatagtgtatgcggcgaccg
agttgctcttgcccggcgtcaatacgggataataccgcgccacatagcagaactttaaaagtgctcatcattggaaaacgttcttcggggcga
aaactctcaaggatcttaccgctgttgagatccagttcgatgtaacccactcgtgcacccaactgatcttcagcatcttttactttcaccagcgttt
ctgggtgagcaaaaacaggaaggcaaaatgccgcaaaaaagggaataagggcgacacggaaatgttgaatactcatactcttcctttttca
atattattgaagcatttatcagggttattgtctcatgagcggatacatatttgaatgtatttagaaaaataaacaaataggggttccgcgcacattt

ccccgaaaagtgccacctgacgtcgacggatcgggagatctcccgatcccctatggtgcactctcagtacaatctgctctgatgccgcatag
ttaagccagtatctgctccctgcttgtgtgttggaggtcgctgagtagtgcgcgagcaaaatttaagctacaacaaggcaaggcttgaccgac
aattgcatgaagaatctgcttagggttaggcgttttgcgctgcttcgcgatgtacgggccagatatacgcgctgacattgattattga

4. **pLV-U6-[empty]-EFSNC-dSaCas9-KRAB-MeCp2-P2A-Puro-WPRE (pCL111) (no gRNA vector)**

gtcgacggatcgggagatctcccgatcccctatggtgcactctcagtacaatctgctctgatgccgcatagttaagccagtatctgctccctgc
ttgtgtgttggaggtcgctgagtagtgcgcgagcaaaatttaagctacaacaaggcaaggcttgaccgacaattgcatgaagaatctgcttag
ggttaggcgttttgcgctgcttcgcgatgtacgggccagatatacgcgttgacattgattattgactagttattaatagtaatcaattacggggtc
attagttcatagcccatatatggagttccgcgttacataacttacggtaaatggcccgcctggctgaccgcccaacgacccccgcccattgac
gtcaataatgacgtatgttcccatagtaacgccaatagggactttccattgacgtcaatgggtggagtatttacggtaaactgcccacttggca
gtacatcaagtgtatcatatgccaagtacgccccctattgacgtcaatgacggtaaatggcccgcctggcattatgcccagtacatgaccttat
gggactttcctacttggcagtacatctacgtattagtcatcgctattaccatggtgatgcggttttggcagtacatcaatgggcgtggatagcgg
tttgactcacggggatttccaagtctccaccccattgacgtcaatgggagtttgttttggcaccaaaatcaacgggactttccaaaatgtcgtaa
caactccgccccattgacgcaaatgggcggtaggcgtgtacggtgggaggtctatataagcagcgcgttttgcctgtactgggtctctctggt
tagaccagatctgagcctgggagctctctggctaactagggaacccactgcttaagcctcaataaagcttgccttgagtgcttcaagtagtgt
gtgcccgtctgttgtgtgactctggtaactagagatccctcagacccttttagtcagtgtggaaaatctctagcagtggcgcccgaacaggga
cttgaaagcgaaagggaaaccagaggagctctctcgacgcaggactcggcttgctgaagcgcgcacggcaagaggcgaggggcggc
gactggtgagtacgccaaaaattttgactagcggaggctagaaggagagagatgggtgcgagagcgtcagtattaagcgggggagaatt
agatcgcgatgggaaaaaattcggttaaggccagggggaaagaaaaaatataaattaaaacatatagtatgggcaagcagggagctagaa
cgattcgcagttaatcctggcctgttagaaacatcagaaggctgtagacaaatactgggacagctacaaccatcccttcagacaggatcaga
agaacttagatcattatataatacagtagcaaccctctattgtgtgcatcaaaggatagagataaaagacaccaaggaagctttagacaagat
agaggaagagcaaaacaaaagtaagaccaccgcacagcaagcggccgctgatcttcagacctggaggaggagatatgagggacaattg
gagaagtgaattatataaatataaagtagtaaaaattgaaccattaggagtagcacccaccaaggcaaagagaagagtggtgcagagaga
aaaaagagcagtgggaataggagctttgttccttgggttcttgggagcagcaggaagcactatgggcgcagcgtcaatgacgctgacggt
acaggccagacaattattgtctggtatagtgcagcagcagaacaatttgctgagggctattgaggcgcaacagcatctgttgcaactcacagt
ctggggcatcaagcagctccaggcaagaatcctggctgtggaaagatacctaaaggatcaacagctcctggggatttggggttgctctgga
aaactcatttgcaccactgctgtgccttggaatgctagttggagtaataaatctctggaacagatttggaatcacacgacctggatggagtggg
acagagaaattaacaattacacaagcttaatacactccttaattgaagaatcgcaaaaccagcaagaaaagaatgaacaagaattattggaat
tagataaatgggcaagtttgtggaattggtttaacataacaaattggctgtggtatataaaattattcataatgatagtaggaggcttggtaggttt
aagaatagtttttgctgtactttctatagtgaatagagttaggcagggatattcaccattatcgtttcagacccacctcccaaccccgaggggac
ccgacaggcccgaaggaatagaagaagaaggtggagagagagacagagacagatccattcgattagtgaacggatcggcactgcgtgc
gccaattctgcagacaaatggcagtattcatccacaattttaaaagaaaaggggggattggggggtacagtgcaggggaaagaatagtaga
cataatagcaacagacatacaaactaaagaattacaaaaacaaattacaaaaattcaaaattttcgggtttattacagggacagcagagatcc
agtttggTTAATTAATGGGCGGGACGTTAACGGGGCGGAACGGTACCgagggcctatttcccatgattc
cttcatatttgcatatacgatacaaggctgttagagagataattagaattaatttgactgtaaacacaaagatattagtacaaaatacgtgacgta
gaaagtaataatttcttgggtagtttgcagttttaaaattatgttttaaaatggactatcatatgcttaccgtaacttgaaagtatttcgatttcttggct
ttatatatcttGTGGAAAGGACGAAAcaccggagacgtgtacacgtctctgTTTtagtactctggaaacagaatctactaaaa
caaggcaaaatgccgtgtttatctcgtcaacttgttggcgagaTTTTTTgaattcgctagctaggtcttgaaaggagtgggaattggctcc

ggtgcccgtcagtgggcagagcgcacatcgcccacagtccccgagaagttggggggaggggtcggcaattgatccggtgcctagagaa
ggtggcgcggggtaaactgggaaagtgatgtcgtgtactggctccgcctttttcccgagggtgggggagaaccgtatataagtgcagtagt
cgccgtgaacgttctttttcgcaacgggtttgccgccagaacacaggaccggtgccaccatgGACTACAAAGACCATGAC
GGTGATTATAAAGATCATGACATaGATTACAAGGATGACGATGACAAGggtggatcgccaaaga
aaaagaggaaagtcgccccaaagaagaagcggaaggtcggtatccacggagtcccagcagccaagcggaactacatcctgggcctgg
ccatcggcatcaccagcgtgggctacggcatcatcgactacgagacacgggacgtgatcgatgccggcgtgcggctgttcaaagaggcc
aacgtggaaaacaacgagggcaggcggagcaagagaggcgccagaaggctgaagcggcggaggcggcatagaatccagagagtga
agaagctgctgttcgactacaacctgctgaccgaccacagcgagctgagcggcatcaacccctacgaggccagagtgaagggcctgagc
cagaagctgagcgaggaagagttctctgccgccctgctgcacctggccaagagaagaggcgtgcacaacgtgaacgaggtggaagag
gacaccggcaacgagctgtccaccaaagagcagatcagccggaacagcaaggccctggaagagaaatacgtggccgaactgcagctg
gaacggctgaagaaagacggcgaagtgcggggcagcatcaacagattcaagaccagcgactacgtgaaagaagccaaacagctgctg
aaggtgcagaaggcctaccaccagctggaccagagcttcatcgacacctacatcgacctgctggaaacccggcggacctactatgaggg
acctggcgagggcagccccttcggctggaaggacatcaaagaatggtacgagatgctgatgggccactgcacctacttccccgaggaact
gcggagcgtgaagtacgcctacaacgccgacctgtacaacgccctgaacgacctgaacaatctcgtgatcaccagggacgagaacgag
aagctggaatattacgagaagttccagatcatcgagaacgtgttcaagcagaagaagaagcccaccctgaagcagatcgccaaagaaatc
ctcgtgaacgaagaggatattaagggctacagagtgaccagcaccggcaagcccgagttcaccaacctgaaggtgtaccacgacatcaa
ggacattaccgcccggaaagagattattgagaacgccgagctgctggatcagattgccaagatcctgaccatctaccagagcagcgagga
catccaggaagaactgaccaatctgaactccgagctgacccaggaagagatcgagcagatctctaatctgaagggctataccggcaccca
caacctgagcctgaaggccatcaacctgatcctggacgagctgtggcacaccaacgacaaccagatcgctatcttcaaccggctgaagct
ggtgcccaagaaggtggacctgtcccagcagaaagagatccccaccaccctggtggacgacttcatcctgagccccgtcgtgaagagaa
gcttcatccagagcatcaaagtgatcaacgccatcatcaagaagtacggcctgcccaacgacatcattatcgagctggcccgcgagaaga
actccaaggacgcccagaaaatgatcaacgagatgcagaagcggaaccggcagaccaacgagcggatcgaggaaatcatccggacca
ccggcaaagagaacgccaagtacctgatcgagaagatcaagctgcacgacatgcaggaaggcaagtgcctgtacagcctggaagccat
ccctctggaagatctgctgaacaaccccttcaactatgaggtggaccacatcatccccagaagcgtgtccttcgacaacagcttcaacaaca
aggtgctcgtgaagcaggaagaagccagcaagaagggcaaccggaccccattccagtacctgagcagcagcgacagcaagatcagct
acgaaaccttcaagaagcacatcctgaatctggccaagggcaagggcagaatcagcaagaccaagaaagagtatctgctggaagaacg
ggacatcaacaggttctccgtgcagaaagacttcatcaaccggaacctggtggataccagatacgccaccagaggcctgatgaacctgct
gcggagctacttcagagtgaacaacctggacgtgaaagtgaagtccatcaatggcggcttcaccagctttctgcggcggaagtggaagttt
aagaaagagcggaacaaggggtacaagcaccacgccgaggacgccctgatcattgccaacgccgatttcatcttcaaagagtggaagaa
actggacaaggccaaaaaagtgatggaaaaccagatgttcgaggaaaagcaggccgagagcatgcccgagatcgaaaccgagcagga
gtacaaagagatcttcatcacccccaccagatcaagcacattaaggacttcaaggactacaagtacagccaccgggtggacaagaagcc
taatagagagctgattaacgacaccctgtactccacccggaaggacgacaagggcaacaccctgatcgtgaacaatctgaacggcctgta
cgacaaggacaatgacaagctgaaaaagctgatcaacaagagccccgaaaagctgctgatgtaccaccacgacccccagacctaccag
aaactgaagctgattatggaacagtacggcgacgagaagaatcccctgtacaagtactacgaggaaaccgggaactacctgaccaagtac
tccaaaaaggacaacggccccgtgatcaagaagattaagtattacggcaacaaactgaacgcccatctggacatcaccgacgactacccc
aacagcagaaacaaggtcgtgaagctgtccctgaagccctacagattcgacgtgtacctggacaatggcgtgtacaagttcgtgaccgtga
agaatctggatgtgatcaaaaaagaaaactactacgaagtgaatagcaagtgctatgaggaagctaagaagctgaagaagatcagcaacc
aggccgagtttatcgcctccttctacaacaacgatctgatcaagatcaacggcgagctgtatagagtgatcggcgtgaacaacgacctgctg
aaccggatcgaagtgaacatgatcgacatcacctaccgcgagtacctggaaaacatgaacgacaagaggcccccccaggatcattaagac
aatcgcctccaagacccagagcattaagaagtacagcacagacattctgggcaacctgtatgaagtgaaatctaagaagcaccctcagatc

atcaaaaagggcaaaaggccggcggccacgaaaaaggcGcTCGAgggtggaggaagtggcgggtcagggtcgggtggcAGC
GCTtcagggtcgggtggcACTAGTcggacactggtgaccttcaaggatgtatttgtggacttcaccagggaggagtggaagctgct
ggacactgctcagcagatcgtgtacagaaatgtgatgctggagaactataagaacctggtttccttgggttatcagcttactaagccagatgtg
atcctccggttggagaagggagaagagccctcgggaggtggttcgggaggtggttcggagggtgtgcaggtgaaaagggtcctggaga
aaagtcctgggaagctccttgtcaagatgccttttcaaacttcgccaggggggcaaggctgaggggggtggggccaccacatccacccagg
tcatggtgatcaaacgccccggcaggaagcgaaaagctgaggccgaccctcaggccattcccaagaaacggggccgaaagccgggga
gtgtggtggcagccgctgccgccgaggccaaaaagaaagccgtgaaggagtcttctatccgatctgtgcaggagacAgtactccccatc
aagaagcgcaagacccgggagGCTAGCggaggtggttcGCCTAGGggaggtggttcgccaaagaagaaacggaaggtgg
gCCGGcccggatccGGCGCAACAAACTTCTCTCTGCTGAAACAAGCCGGAGATGTCGAAG
AGAATCCTGGACCGACCGAGTACAAGCCCACGGTGCGCCTCGCCACCCGCGACGAC
GTCCCCAGGGCCGTACGCACCCTCGCCGCCGCGTTCGCCGACTACCCCGCCACGCGC
CACACCGTCGATCCGGACCGCCACATCGAGCGGGTCACCGAGCTGCAAGAACTCTTC
CTCACGCGCGTCGGGCTCGACATCGGCAAGGTGTGGGTCGCGGACGACGGCGCCGC
GGTGGCGGTCTGGACCACGCCGGAGAGCGTCGAAGCGGGGGCGGTGTTCGCCGAGA
TCGGCCCGCGCATGGCCGAGTTGAGCGGTTCCCGGCTGGCCGCGCAGCAACAGATG
GAAGGCCTCCTGGCGCCGCACCGGCCCAAGGAGCCCGCGTGGTTCCTGGCCACCGT
CGGAGTCTCGCCCGACCACCAGGGCAAGGGTCTGGGCAGCGCCGTCGTGCTCCCCG
GAGTGGAGGCGGCCGAGCGCGCCGGGGTGCCCGCCTTCCTGGAGACCTCCGCGCCC
CGCAACCTCCCCTTCTACGAGCGGCTCGGCTTCACCGTCACCGCCGACGTCGAGGTG
CCCGAAGGACCGCGCACCTGGTGCATGACCCGCAAGCCCGGTGCCTGAACGCGTTA
AGTCGACAATCAACCTCTGGATTACAAAATTTGTGAAAGATTGACTGGTATTCTTAAC
TATGTTGCTCCTTTTACGCTATGTGGATACGCTGCTTTAATGCCTTTGTATCATGCTATT
GCTTCCCGTATGGCTTTCATTTTCTCCTCCTTGTATAAATCCTGGTTGCTGTCTCTTTAT
GAGGAGTTGTGGCCCGTTGTCAGGCAACGTGGCGTGGTGTGCACTGTGTTTGCTGAC
GCAACCCCCACTGGTTGGGGCATTGCCACCACCTGTCAGCTCCTTTCCGGGACTTTC
GCTTTCCCCCTCCCTATTGCCACGGCGGAACTCATCGCCGCCTGCCTTGCCCGCTGCT
GGACAGGGGCTCGGCTGTTGGGCACTGACAATTCCGTGGTGTTGTCGGGGAAATCAT
CGTCCTTTCCTTGGCTGCTCGCCTGTGTTGCCACCTGGATTCTGCGCGGGACGTCCTT
CTGCTACGTCCCTTCGGCCCTCAATCCAGCGGACCTTCCTTCCCGCGGCCTGCTGCCG
GCTCTGCGGCCTCTTCCGCGTCTTCGCCTTCGCCCTCAGACGAGTCGGATCTCCCTTT
GGGCCGCCTCCCCGCGTCGACTTTAAGACCAATGACTTACAAGGCAGCTGTAGATCTT
AGCCACTTTTTAAAAGAAAAGGGGGGACTGGAAGGGCTAATTCACTCCCAACGAAG
ACAAGATCTGCTTTTTGCTTGTACTGGGTCTCTCTGGTTAGACCAGATCTGAGCCTGG
GAGCTCTCTGGCTAACTAGGGAACCCACTGCTTAAGCCTCAATAAAGCTTGCCTTGA
GTGCTTCAAGTAGTGTGTGCCCGTCTGTTGTGTGACTCTGGTAACTAGAGATCCCTCA
GACCCTTTTAGTCAGTGTGGAAAATCTCTAGCAGggcccgtttaaacccgctgatcagcctcgactgtgcctt
ctagttgccagccatctgttgtttgcccctcccccgtgccttccttgaccctggaaggtgccactcccactgtcctttcctaataaaatgaggaa
attgcatcgcattgtctgagtaggtgtcattctattctggggggtggggtggggcaggacagcaagggggaggattgggaagacaatagca
ggcatgctggggatgcggtgggctctatggcttctgaggcggaaagaaccagctggggctctagggggtatccccacgcgccctgtagc
ggcgcattaagcgcggcgggtgtggtggttacgcgcagcgtgaccgctacacttgccagcgccctagcgcccgctcctttcgctttcttccc

ttcctttctcgccacgttcgccggctttccccgtcaagctctaaatcgggggctccctttagggttccgatttagtgctttacggcacctcgaccc
caaaaaacttgattagggtgatggttcacgtagtgggccatcgccctgatagacggtttttcgccctttgacgttggagtccacgttctttaata
gtggactcttgttccaaactggaacaacactcaaccctatctcggtctattcttttgatttataatggttacaaataaagcaatagcatcacaaattt
cacaaataaagcatttttttcactgcattctagttgtggtttgtccaaactcatcaatgtatcttatcatgtctgtataccgtcgacctctagctagag
cttggcgtaatcatggtcatagctgtttcctgtgtgaaattgttatccgctcacaattccacacaacatacgagccggaagcataaagtgtaaag
cctggggtgcctaatgagtgagctaactcacattaattgcgttgcgctcactgcccgctttccagtcgggaaacctgtcgtgccagctgcatta
atgaatcggccaacgcgcggggagaggcggtttgcgtattgggcgctcttccgcttcctcgctcactgactcgctgcgctcggtcgttcggc
tgcggcgagcggtatcagctcactcaaaggcggtaatacggttatccacagaatcaggggataacgcaggaaagaacatgtgagcaaaa
ggccagcaaaaggccaggaaccgtaaaaaggccgcgttgctggcgtttttccataggctccgcccccctgacgagcatcacaaaaatcga
cgctcaagtcagaggtggcgaaacccgacaggactataaagataccaggcgtttccccctggaagctccctcgtgcgctctcctgttccga
ccctgccgcttaccggatacctgtccgcctttctcccttcgggaagcgtggcgctttctcatagctcacgctgtaggtatctcagttcggtgtag
gtcgttcgctccaagctgggctgtgtgcacgaaccccccgttcagcccgaccgctgcgccttatccggtaactatcgtcttgagtccaaccc
ggtaagacacgacttatcgccactggcagcagccactggtaacaggattagcagagcgaggtatgtaggcggtgctacagagttcttgaa
gtggtggcctaactacggctacactagaagaacagtatttggtatctgcgctctgctgaagccagttaccttcggaaaaagagttggtagctct
tgatccggcaaacaaaccaccgctggtagcggtggtttttttgtttgcaagcagcagattacgcgcagaaaaaaaggatctcaagaagatcc
tttgatcttttctacggggtctgacgctcagtggaacgaaaactcacgttaagggattttggtcatgagattatcaaaaaggatcttcacctagat
ccttttaaattaaaaatgaagttttaaatcaatctaaagtatatatgagtaaacttggtctgacagttaccaatgcttaatcagtgaggcacctatct
cagcgatctgtctatttcgttcatccatagttgcctgactccccgtcgtgtagataactacgatacgggagggcttaccatctggccccagtgct
gcaatgataccgcgagacccacgctcaccggctccagatttatcagcaataaaccagccagccggaagggccgagcgcagaagtggtc
ctgcaactttatccgcctccatccagtctattaattgttgccgggaagctagagtaagtagttcgccagttaatagtttgcgcaacgttgttgcca
ttgctacaggcatcgtggtgtcacgctcgtcgtttggtatggcttcattcagctccggttcccaacgatcaaggcgagttacatgatcccccat
gttgtgcaaaaaagcggttagctccttcggtcctccgatcgttgtcagaagtaagttggccgcagtgttatcactcatggttatggcagcactg
cataattctcttactgtcatgccatccgtaagatgcttttctgtgactggtgagtactcaaccaagtcattctgagaatagtgtatgcggcgaccg
agttgctcttgcccggcgtcaatacgggataataccgcgccacatagcagaactttaaaagtgctcatcattggaaaacgttcttcggggcga
aaactctcaaggatcttaccgctgttgagatccagttcgatgtaacccactcgtgcacccaactgatcttcagcatcttttactttcaccagcgttt
ctgggtgagcaaaaacaggaaggcaaaatgccgcaaaaaagggaataagggcgacacggaaatgttgaatactcatactcttcctttttca
atattattgaagcatttatcagggttattgtctcatgagcggatacatatttgaatgtatttagaaaaataaacaaataggggttccgcgcacattt
ccccgaaaagtgccacctgac

5. **pBK1841- pLV-U6-[SNCAintron1 gRNA 1]-EFSNC-dSaCas9-KRAB-MeCp2-P2A-Puro-WPRE (THERAPEUTIC VECTOR)**

gtcgacggatcgggagatctcccgatcccctatggtgcactctcagtacaatctgctctgatgccgcatagttaagccagtatctgctccctgc
ttgtgtgttggaggtcgctgagtagtgcgcgagcaaaatttaagctacaacaaggcaaggcttgaccgacaattgcatgaagaatctgcttag
ggttaggcgttttgcgctgcttcgcgatgtacgggccagatatacgcgttgacattgattattgactagttattaatagtaatcaattacggggtc
attagttcatagcccatatatggagttccgcgttacataacttacggtaaatggcccgcctggctgaccgcccaacgacccccgcccattgac
gtcaataatgacgtatgttcccatagtaacgccaatagggactttccattgacgtcaatgggtggagtatttacggtaaactgcccacttggca
gtacatcaagtgtatcatatgccaagtacgccccctattgacgtcaatgacggtaaatggcccgcctggcattatgcccagtacatgaccttat
gggactttcctacttggcagtacatctacgtattagtcatcgctattaccatggtgatgcggttttggcagtacatcaatgggcgtggatagcgg
tttgactcacggggatttccaagtctccaccccattgacgtcaatgggagtttgttttggcaccaaaatcaacgggactttccaaaatgtcgtaa
caactccgccccattgacgcaaatgggcggtaggcgtgtacggtgggaggtctatataagcagcgcgttttgcctgtactgggtctctctggt

tagaccagatctgagcctgggagctctctggctaactagggaacccactgcttaagcctcaataaagcttgccttgagtgcttcaagtagtgt
gtgcccgtctgttgtgtgactctggtaactagagatccctcagacccttttagtcagtgtggaaaatctctagcagtggcgcccgaacaggga
cttgaaagcgaaagggaaaccagaggagctctctcgacgcaggactcggcttgctgaagcgcgcacggcaagaggcgaggggcggc
gactggtgagtacgccaaaaattttgactagcggaggctagaaggagagagatgggtgcgagagcgtcagtattaagcgggggagaatt
agatcgcgatgggaaaaaattcggttaaggccagggggaaagaaaaaatataaattaaaacatatagtatgggcaagcagggagctagaa
cgattcgcagttaatcctggcctgttagaaacatcagaaggctgtagacaaatactgggacagctacaaccatcccttcagacaggatcaga
agaacttagatcattatataatacagtagcaaccctctattgtgtgcatcaaaggatagagataaaagacaccaaggaagctttagacaagat
agaggaagagcaaaacaaaagtaagaccaccgcacagcaagcggccgctgatcttcagacctggaggaggagatatgagggacaattg
gagaagtgaattatataaatataaagtagtaaaaattgaaccattaggagtagcacccaccaaggcaaagagaagagtggtgcagagaga
aaaaagagcagtgggaataggagctttgttccttgggttcttgggagcagcaggaagcactatgggcgcagcgtcaatgacgctgacggt
acaggccagacaattattgtctggtatagtgcagcagcagaacaatttgctgagggctattgaggcgcaacagcatctgttgcaactcacagt
ctggggcatcaagcagctccaggcaagaatcctggctgtggaaagatacctaaaggatcaacagctcctggggatttggggttgctctgga
aaactcatttgcaccactgctgtgccttggaatgctagttggagtaataaatctctggaacagatttggaatcacacgacctggatggagtggg
acagagaaattaacaattacacaagcttaatacactccttaattgaagaatcgcaaaaccagcaagaaaagaatgaacaagaattattggaat
tagataaatgggcaagtttgtggaattggtttaacataacaaattggctgtggtatataaaattattcataatgatagtaggaggcttggtaggttt
aagaatagtttttgctgtactttctatagtgaatagagttaggcagggatattcaccattatcgtttcagacccacctcccaaccccgaggggac
ccgacaggcccgaaggaatagaagaagaaggtggagagagagacagagacagatccattcgattagtgaacggatcggcactgcgtgc
gccaattctgcagacaaatggcagtattcatccacaattttaaaagaaaaggggggattggggggtacagtgcaggggaaagaatagtaga
cataatagcaacagacatacaaactaaagaattacaaaaacaaattacaaaaattcaaaattttcgggtttattacagggacagcagagatcc
agtttggTTAATTAATGGGCGGACGTTAACGGGGCGGAACGGTACCgagggcctatttcccatgattc
cttcatatttgcatatacgatacaaggctgttagagagataattagaattaatttgactgtaaacacaaagatattagtacaaaatacgtgacgta
gaaagtaataatttcttgggtagtttgcagttttaaaattatgttttaaaatggactatcatatgcttaccgtaacttgaaagtatttcgatttcttggct
ttatatatcttGTGGAAAGGACGAAAcaccgACCTCCCAGAGACCTGGCCCAgTTTtagtactctggaa
acagaatctactaaaacaaggcaaaatgccgtgtttatctcgtcaacttgttggcgagaTTTTTTgaattcgctagctaggtcttgaaagg
agtgggaattggctccggtgcccgtcagtgggcagagcgcacatcgcccacagtccccgagaagttggggggaggggtcggcaattgat
ccggtgcctagagaaggtggcgcggggtaaactgggaaagtgatgtcgtgtactggctccgcctttttcccgagggtgggggagaaccgt
atataagtgcagtagtcgccgtgaacgttctttttcgcaacgggtttgccgccagaacacaggaccggtgccaccatgGACTACAA
AGACCATGACGGTGATTATAAAGATCATGACATaGATTACAAGGATGACGATGACAAG
ggtggatcgccaaagaaaaagaggaaagtcgccccaaagaagaagcggaaggtcggtatccacggagtcccagcagccaagcggaa
ctacatcctgggcctggccatcggcatcaccagcgtgggctacggcatcatcgactacgagacacgggacgtgatcgatgccggcgtgc
ggctgttcaaagaggccaacgtggaaaacaacgagggcaggcggagcaagagaggcgccagaaggctgaagcggcggaggcggca
tagaatccagagagtgaagaagctgctgttcgactacaacctgctgaccgaccacagcgagctgagcggcatcaacccctacgaggcca
gagtgaagggcctgagccagaagctgagcgaggaagagttctctgccgccctgctgcacctggccaagagaagaggcgtgcacaacgt
gaacgaggtggaagaggacaccggcaacgagctgtccaccaaagagcagatcagccggaacagcaaggccctggaagagaaatacg
tggccgaactgcagctggaacggctgaagaaagacggcgaagtgcggggcagcatcaacagattcaagaccagcgactacgtgaaag
aagccaaacagctgctgaaggtgcagaaggcctaccaccagctggaccagagcttcatcgacacctacatcgacctgctggaaacccgg
cggacctactatgagggacctggcgagggcagccccttcggctggaaggacatcaaagaatggtacgagatgctgatgggccactgcac
ctacttccccgaggaactgcggagcgtgaagtacgcctacaacgccgacctgtacaacgccctgaacgacctgaacaatctcgtgatcac
cagggacgagaacgagaagctggaatattacgagaagttccagatcatcgagaacgtgttcaagcagaagaagaagcccaccctgaagc
agatcgccaaagaaatcctcgtgaacgaagaggatattaagggctacagagtgaccagcaccggcaagcccgagttcaccaacctgaag

gtgtaccacgacatcaaggacattaccgcccggaaagagattattgagaacgccgagctgctggatcagattgccaagatcctgaccatct
accagagcagcgaggacatccaggaagaactgaccaatctgaactccgagctgacccaggaagagatcgagcagatctctaatctgaag
ggctataccggcacccacaacctgagcctgaaggccatcaacctgatcctggacgagctgtggcacaccaacgacaaccagatcgctatc
ttcaaccggctgaagctggtgcccaagaaggtggacctgtcccagcagaaagagatccccaccaccctggtggacgacttcatcctgagc
cccgtcgtgaagagaagcttcatccagagcatcaaagtgatcaacgccatcatcaagaagtacggcctgcccaacgacatcattatcgagc
tggcccgcgagaagaactccaaggacgcccagaaaatgatcaacgagatgcagaagcggaaccggcagaccaacgagcggatcgag
gaaatcatccggaccaccggcaaagagaacgccaagtacctgatcgagaagatcaagctgcacgacatgcaggaaggcaagtgcctgt
acagcctggaagccatccctctggaagatctgctgaacaaccccttcaactatgaggtggaccacatcatccccagaagcgtgtccttcgac
aacagcttcaacaacaaggtgctcgtgaagcaggaagaagccagcaagaagggcaaccggaccccattccagtacctgagcagcagcg
acagcaagatcagctacgaaaccttcaagaagcacatcctgaatctggccaagggcaagggcagaatcagcaagaccaagaaagagtat
ctgctggaagaacgggacatcaacaggttctccgtgcagaaagacttcatcaaccggaacctggtggataccagatacgccaccagaggc
ctgatgaacctgctgcggagctacttcagagtgaacaacctggacgtgaaagtgaagtccatcaatggcggcttcaccagctttctgcggcg
gaagtggaagtttaagaaagagcggaacaaggggtacaagcaccacgccgaggacgccctgatcattgccaacgccgatttcatcttcaa
agagtggaagaaactggacaaggccaaaaaagtgatggaaaaccagatgttcgaggaaaagcaggccgagagcatgcccgagatcga
aaccgagcaggagtacaaagagatcttcatcacccccccaccagatcaagcacattaaggacttcaaggactacaagtacagccaccgggt
ggacaagaagcctaatagagagctgattaacgacaccctgtactccacccggaaggacgacaagggcaacaccctgatcgtgaacaatct
gaacggcctgtacgacaaggacaatgacaagctgaaaaagctgatcaacaagagccccgaaaagctgctgatgtaccaccacgacccccc
agacctaccagaaactgaagctgattatggaacagtacggcgacgagaagaatcccctgtacaagtactacgaggaaaccgggaactac
ctgaccaagtactccaaaaaggacaacggccccgtgatcaagaagattaagtattacggcaacaaactgaacgcccatctggacatcacc
gacgactaccccaacagcagaaacaaggtcgtgaagctgtccctgaagccctacagattcgacgtgtacctggacaatggcgtgtacaag
ttcgtgaccgtgaagaatctggatgtgatcaaaaaagaaaactactacgaagtgaatagcaagtgctatgaggaagctaagaagctgaaga
agatcagcaaccaggccgagtttatcgcctccttctacaacaacgatctgatcaagatcaacggcgagctgtatagagtgatcggcgtgaac
aacgacctgctgaaccggatcgaagtgaacatgatcgacatcacctaccgcgagtacctggaaaacatgaacgacaagaggcccccccag
gatcattaagacaatcgcctccaagacccagagcattaagaagtacagcacagacattctgggcaacctgtatgaagtgaaatctaagaag
caccctcagatcatcaaaaagggcaaaaggccggcggccacgaaaaaggcGcTCGAgggtggaggaagtggcgggtcagggtc
gggtggcAGCGCTtcagggtcgggtggcACTAGTcggacactggtgaccttcaaggatgtatttgtggacttcaccagggagga
gtggaagctgctggacactgctcagcagatcgtgtacagaaatgtgatgctggagaactataagaacctggtttccttgggttatcagcttact
aagccagatgtgatcctccggttggagaagggagaagagccctcgggaggtggttcgggaggtggttcggagggtgtgcaggtgaaaa
gggtcctggagaaaagtcctgggaagctccttgtcaagatgccttttcaaacttcgccagggggcaaggctgagggggggtggggccacca
catccacccaggtcatggtgatcaaacgccccggcaggaagcgaaaagctgaggccgaccctcaggccattcccaagaaacggggccg
aaagccggggagtgtggtggcagccgctgccgccgaggccaaaaagaaagccgtgaaggagtcttctatccgatctgtgcaggagacA
gtactccccatcaagaagcgcaagacccgggagGCTAGCggaggtggttcGCCTAGGggaggtggttcgccaaagaagaaa
cggaaggtgggCCGGcccggatccGGCGCAACAAACTTCTCTCTGCTGAAACAAGCCGGAGAT
GTCGAAGAGAATCCTGGACCGACCGAGTACAAGCCCACGGTGCGCCTCGCCACCCG
CGACGACGTCCCCAGGGCCGTACGCACCCTCGCCGCCGCGTTCGCCGACTACCCCGC
CACGCGCCACACCGTCGATCCGGACCGCCACATCGAGCGGGTCACCGAGCTGCAAG
AACTCTTCCTCACGCGCGTCGGGCTCGACATCGGCAAGGTGTGGGTCGCGGACGACG
GCGCCGCGGTGGCGGTCTGGACCACGCCGGAGAGCGTCGAAGCGGGGGCGGTGTTC
GCCGAGATCGGCCCGCGCATGGCCGAGTTGAGCGGTTCCCGGCTGGCCGCGCAGCA
ACAGATGGAAGGCCTCCTGGCGCCGCACCGGCCCAAGGAGCCCGCGTGGTTCCTGG

CCACCGTCGGAGTCTCGCCCGACCACCAGGGCAAGGGTCTGGGCAGCGCCGTCGTG
CTCCCCGGAGTGGAGGCGGCCGAGCGCGCCGGGGTGCCCGCCTTCCTGGAGACCTC
CGCGCCCCGCAACCTCCCCTTCTACGAGCGGCTCGGCTTCACCGTCACCGCCGACGT
CGAGGTGCCCGAAGGACCGCGCACCTGGTGCATGACCCGCAAGCCCGGTGCCTGAA
CGCGTTAAGTCGACAATCAACCTCTGGATTACAAAATTTGTGAAAGATTGACTGGTAT
TCTTAACTATGTTGCTCCTTTTACGCTATGTGGATACGCTGCTTTAATGCCTTTGTATCA
TGCTATTGCTTCCCGTATGGCTTTCATTTTCTCCTCCTTGTATAAATCCTGGTTGCTGTC
TCTTTATGAGGAGTTGTGGCCCGTTGTCAGGCAACGTGGCGTGGTGTGCACTGTGTTT
GCTGACGCAACCCCCACTGGTTGGGGCATTGCCACCACCTGTCAGCTCCTTTCCGGG
ACTTTCGCTTTCCCCCTCCCTATTGCCACGGCGGAACTCATCGCCGCCTGCCTTGCCC
GCTGCTGGACAGGGGCTCGGCTGTTGGGCACTGACAATTCCGTGGTGTTGTCGGGGA
AATCATCGTCCTTTCCTTGGCTGCTCGCCTGTGTTGCCACCTGGATTCTGCGCGGGAC
GTCCTTCTGCTACGTCCCTTCGGCCCTCAATCCAGCGGACCTTCCTTCCCGCGGCCTG
CTGCCGGCTCTGCGGCCTCTTCCGCGTCTTCGCCTTCGCCCTCAGACGAGTCGGATCT
CCCTTTGGGCCGCCTCCCCGCGTCGACTTTAAGACCAATGACTTACAAGGCAGCTGTA
GATCTTAGCCACTTTTTAAAAGAAAAGGGGGGACTGGAAGGGCTAATTCACTCCCAA
CGAAGACAAGATCTGCTTTTTGCTTGTACTGGGTCTCTCTGGTTAGACCAGATCTGAG
CCTGGGAGCTCTCTGGCTAACTAGGGAACCCACTGCTTAAGCCTCAATAAAGCTTGC
CTTGAGTGCTTCAAGTAGTGTGTGCCCGTCTGTTGTGTGACTCTGGTAACTAGAGATC
CCTCAGACCCTTTTAGTCAGTGTGGAAAATCTCTAGCAGggcccgtttaaacccgctgatcagcctcga
ctgtgccttctagttgccagccatctgttgtttgcccctcccccgtgccttccttgaccctggaaggtgccactcccactgtcctttcctaataaa
atgaggaaattgcatcgcattgtctgagtaggtgtcattctattctggggggtggggtggggcaggacagcaagggggaggattgggaag
acaatagcaggcatgctggggatgcggtgggctctatggcttctgaggcggaaagaaccagctggggctctagggggtatccccacgcg
ccctgtagcggcgcattaagcgcggcgggtgtggtggttacgcgcagcgtgaccgctacacttgccagcgccctagcgcccgctcctttc
gctttcttcccttcctttctcgccacgttcgccggctttccccgtcaagctctaaatcgggggctccctttagggttccgatttagtgctttacggc
acctcgaccccaaaaaacttgattagggtgatggttcacgtagtgggccatcgccctgatagacggtttttcgccctttgacgttggagtccac
gttctttaatagtggactcttgttccaaactggaacaacactcaaccctatctcggtctattcttttgatttataatggttacaaataaagcaatagc
atcacaaatttcacaaataaagcatttttttcactgcattctagttgtggtttgtccaaactcatcaatgtatcttatcatgtctgtataccgtcgacct
ctagctagagcttggcgtaatcatggtcatagctgtttcctgtgtgaaattgttatccgctcacaattccacacaacatacgagccggaagcata
aagtgtaaagcctggggtgcctaatgagtgagctaactcacattaattgcgttgcgctcactgcccgctttccagtcgggaaacctgtcgtgc
cagctgcattaatgaatcggccaacgcgcggggagaggcggtttgcgtattgggcgctcttccgcttcctcgctcactgactcgctgcgctc
ggtcgttcggctgcggcgagcggtatcagctcactcaaaggcggtaatacggttatccacagaatcaggggataacgcaggaaagaacat
gtgagcaaaaggccagcaaaaggccaggaaccgtaaaaaggccgcgttgctggcgtttttccataggctccgcccccctgacgagcatc
acaaaaatcgacgctcaagtcagaggtggcgaaacccgacaggactataaagataccaggcgtttccccctggaagctccctcgtgcgct
ctcctgttccgaccctgccgcttaccggatacctgtccgcctttctcccttcgggaagcgtggcgctttctcatagctcacgctgtaggtatctc
agttcggtgtaggtcgttcgctccaagctgggctgtgtgcacgaaccccccgttcagcccgaccgctgcgccttatccggtaactatcgtctt
gagtccaacccggtaagacacgacttatcgccactggcagcagccactggtaacaggattagcagagcgaggtatgtaggcggtgctaca
gagttcttgaagtggtggcctaactacggctacactagaagaacagtatttggtatctgcgctctgctgaagccagttaccttcggaaaaaga
gttggtagctcttgatccggcaaacaaaccaccgctggtagcggtggtttttttgtttgcaagcagcagattacgcgcagaaaaaaaggatct
caagaagatcctttgatcttttctacggggtctgacgctcagtggaacgaaaactcacgttaagggattttggtcatgagattatcaaaaaggat

cttcacctagatccttttaaattaaaaatgaagttttaaatcaatctaaagtatatatgagtaaacttggtctgacagttaccaatgcttaatcagtg
aggcacctatctcagcgatctgtctatttcgttcatccatagttgcctgactccccgtcgtgtagataactacgatacgggagggcttaccatct
ggccccagtgctgcaatgataccgcgagacccacgctcaccggctccagatttatcagcaataaaccagccagccggaagggccgagc
gcagaagtggtcctgcaactttatccgcctccatccagtctattaattgttgccgggaagctagagtaagtagttcgccagttaatagtttgcgc
aacgttgttgccattgctacaggcatcgtggtgtcacgctcgtcgtttggtatggcttcattcagctccggttcccaacgatcaaggcgagttac
atgatcccccatgttgtgcaaaaaagcggttagctccttcggtcctccgatcgttgtcagaagtaagttggccgcagtgttatcactcatggtta
tggcagcactgcataattctcttactgtcatgccatccgtaagatgcttttctgtgactggtgagtactcaaccaagtcattctgagaatagtgtat
gcggcgaccgagttgctcttgcccggcgtcaatacgggataataccgcgccacatagcagaactttaaaagtgctcatcattggaaaacgtt
cttcggggcgaaaactctcaaggatcttaccgctgttgagatccagttcgatgtaacccactcgtgcacccaactgatcttcagcatcttttactt
tcaccagcgtttctgggtgagcaaaaacaggaaggcaaaatgccgcaaaaaagggaataagggcgacacggaaatgttgaatactcatac
tcttcctttttcaatattattgaagcatttatcagggttattgtctcatgagcggatacatatttgaatgtatttagaaaaataaacaaataggggttc
cgcgcacatttccccgaaaagtgccacctgac

### 6. pCL223- pLV-U6-[hSNCA gRNA 2]-EFSNC-dSpCas9-DNMT3a-P2A-PURO-WPRE (THERAPUTIC VECTOR)

ctagttattaatagtaatcaattacggggtcattagttcatagcccatatatggagttccgcgttacataacttacggtaaatggcccgcctggct
gaccgcccaacgacccccgcccattgacgtcaataatgacgtatgttcccatagtaacgccaatagggactttccattgacgtcaatgggtg
gagtatttacggtaaactgcccacttggcagtacatcaagtgtatcatatgccaagtacgccccctattgacgtcaatgacggtaaatggccc
gcctggcattatgcccagtacatgaccttatgggactttcctacttggcagtacatctacgtattagtcatcgctattaccatggtgatgcggtttt
ggcagtacatcaatgggcgtggatagcggtttgactcacggggatttccaagtctccaccccattgacgtcaatgggagtttgttttggcacc
aaaatcaacgggactttccaaaatgtcgtaacaactccgccccattgacgcaaatgggcggtaggcgtgtacggtgggaggtctatataag
cagcgcgttttgcctgtactgggtctctctggttagaccagatctgagcctgggagctctctggctaactagggaacccactgcttaagcctca
ataaagcttgccttgagtgcttcaagtagtgtgtgcccgtctgttgtgtgactctggtaactagagatccctcagacccttttagtcagtgtggaa
aatctctagcagtggcgcccgaacagggacttgaaagcgaaagggaaaccagaggagctctctcgacgcaggactcggcttgctgaagc
gcgcacggcaagaggcgaggggcggcgactggtgagtacgccaaaaattttgactagcggaggctagaaggagagagatgggtgcga
gagcgtcagtattaagcgggggagaattagatcgcgatgggaaaaaattcggttaaggccagggggaaagaaaaaatataaattaaaaca
tatagtatgggcaagcagggagctagaacgattcgcagttaatcctggcctgttagaaacatcagaaggctgtagacaaatactgggacag
ctacaaccatcccttcagacaggatcagaagaacttagatcattatataatacagtagcaaccctctattgtgtgcatcaaaggatagagataa
aagacaccaaggaagctttagacaagatagaggaagagcaaaacaaaagtaagaccaccgcacagcaagcggccgctgatcttcagac
ctggaggaggagatatgagggacaattggagaagtgaattatataaatataaagtagtaaaaattgaaccattaggagtagcacccaccaa
ggcaaagagaagagtggtgcagagagaaaaaagagcagtgggaataggagctttgttccttgggttcttgggagcagcaggaagcactat
gggcgcagcgtcaatgacgctgacggtacaggccagacaattattgtctggtatagtgcagcagcagaacaatttgctgagggctattgag
gcgcaacagcatctgttgcaactcacagtctggggcatcaagcagctccaggcaagaatcctggctgtggaaagatacctaaaggatcaa
cagctcctggggatttggggttgctctggaaaactcatttgcaccactgctgtgccttggaatgctagttggagtaataaatctctggaacagat
ttggaatcacacgacctggatggagtgggacagagaaattaacaattacacaagcttaatacactccttaattgaagaatcgcaaaaccagc
aagaaaagaatgaacaagaattattggaattagataaatgggcaagtttgtggaattggtttaacataacaaattggctgtggtatataaaattat
tcataatgatagtaggaggcttggtaggtttaagaatagtttttgctgtactttctatagtgaatagagttaggcagggatattcaccattatcgttt
cagacccacctcccaaccccgaggggacccgacaggcccgaaggaatagaagaagaaggtggagagagagacagagacagatccat
tcgattagtgaacggatcggcactgcgtgcgccaattctgcagacaaatggcagtattcatccacaattttaaaagaaaaggggggattggg
gggtacagtgcaggggaaagaatagtagacataatagcaacagacatacaaactaaagaattacaaaaacaaattacaaaaattcaaaattt
tcgggtttattacagggacagcagagatccagtttggTTAATTAATGGGCGGGACGTTAACGGGGCGGAAC
GGTACCgagggcctatttcccatgattccttcatatttgcatatacgatacaaggctgttagagagataattagaattaatttgactgtaaac
acaaagatattagtacaaaatacgtgacgtagaaagtaataatttcttgggtagtttgcagttttaaaattatgttttaaaatggactatcatatgctt

accgtaacttgaaagtatttcgatttcttggctttatatatcttGTGGAAAGGACGAAAcaccgCTGCTCAGGGTAG
ATAGCTGgTTTtagtactctggaaacagaatctactaaaacaaggcaaaatgccgtgtttatctcgtcaacttgttggcgagaTTT
TTTgaattcgctagctaggtcttgaaaggagtgggaattggctccggtgcccgtcagtgggcagagcgcacatcgcccacagtccccga
gaagttggggggaggggtcggcaattgatccggtgcctagagaaggtggcgcggggtaaactgggaaagtgatgtcgtgtactggctcc
gcctttttcccgagggtgggggagaaccgtatataagtgcagtagtcgccgtgaacgttctttttcgcaacgggtttgccgccagaacacag
gaccggtgccaccatgGACTACAAAGACCATGACGGTGATTATAAAGATCATGACATaGATT
ACAAGGATGACGATGACAAGggtggatcgccaaagaaaaagaggaaagtcgccccaaagaagaagcggaaggtcg
gtatccacggagtcccagcagccaagcggaactacatcctgggcctggccatcggcatcaccagcgtgggctacggcatcatcgactac
gagacacgggacgtgatcgatgccggcgtgcggctgttcaaagaggccaacgtggaaaacaacgagggcaggcggagcaagagagg
cgccagaaggctgaagcggcggaggcggcatagaatccagagagtgaagaagctgctgttcgactacaacctgctgaccgaccacagc
gagctgagcggcatcaacccctacgaggccagagtgaagggcctgagccagaagctgagcgaggaagagttctctgccgccctgctgc
acctggccaagagaagaggcgtgcacaacgtgaacgaggtggaagaggacaccggcaacgagctgtccaccaaagagcagatcagc
cggaacagcaaggccctggaagagaaatacgtggccgaactgcagctggaacggctgaagaaagacggcgaagtgcggggcagcat
caacagattcaagaccagcgactacgtgaaagaagccaaacagctgctgaaggtgcagaaggcctaccaccagctggaccagagcttca
tcgacacctacatcgacctgctggaaacccggcggacctactatgagggacctggcgagggcagccccttcggctggaaggacatcaaa
gaatggtacgagatgctgatgggccactgcacctacttccccgaggaactgcggagcgtgaagtacgcctacaacgccgacctgtacaac
gccctgaacgacctgaacaatctcgtgatcaccagggacgagaacgagaagctggaatattacgagaagttccagatcatcgagaacgtg
ttcaagcagaagaagaagcccaccctgaagcagatcgccaaagaaatcctcgtgaacgaagaggatattaagggctacagagtgaccag
caccggcaagcccgagttcaccaacctgaaggtgtaccacgacatcaaggacattaccgcccggaaagagattattgagaacgccgagc
tgctggatcagattgccaagatcctgaccatctaccagagcagcgaggacatccaggaagaactgaccaatctgaactccgagctgaccc
aggaagagatcgagcagatctctaatctgaagggctataccggcacccacaacctgagcctgaaggccatcaacctgatcctggacgagc
tgtggcacaccaacgacaaccagatcgctatcttcaaccggctgaagctggtgcccaagaaggtggacctgtcccagcagaaagagatcc
ccaccaccctggtggacgacttcatcctgagccccgtcgtgaagagaagcttcatccagagcatcaaagtgatcaacgccatcatcaagaa
gtacggcctgcccaacgacatcattatcgagctggcccgcgagaagaactccaaggacgcccagaaaatgatcaacgagatgcagaagc
ggaaccggcagaccaacgagcggatcgaggaaatcatccggaccaccggcaaagagaacgccaagtacctgatcgagaagatcaagc
tgcacgacatgcaggaaggcaagtgcctgtacagcctggaagccatccctctggaagatctgctgaacaaccccttcaactatgaggtgga
ccacatcatccccagaagcgtgtccttcgacaacagcttcaacaacaaggtgctcgtgaagcaggaagaagccagcaagaagggcaacc
ggaccccattccagtacctgagcagcagcgacagcaagatcagctacgaaaccttcaagaagcacatcctgaatctggccaagggcaag
ggcagaatcagcaagaccaagaaagagtatctgctggaagaacgggacatcaacaggttctccgtgcagaaagacttcatcaaccggaa
cctggtggataccagatacgccaccagaggcctgatgaacctgctgcggagctacttcagagtgaacaacctggacgtgaaagtgaagtc
catcaatggcggcttcaccagctttctgcggcggaagtggaagtttaagaaagagcggaacaaggggtacaagcaccacgccgaggacg
ccctgatcattgccaacgccgatttcatcttcaaagagtggaagaaactggacaaggccaaaaaagtgatggaaaaccagatgttcgagga
aaagcaggccgagagcatgcccgagatcgaaaccgagcaggagtacaaagagatcttcatcacccccccaccagatcaagcacattaag
gacttcaaggactacaagtacagccaccgggtggacaagaagcctaatagagagctgattaacgacaccctgtactccacccggaagga
cgacaagggcaacaccctgatcgtgaacaatctgaacggcctgtacgacaaggacaatgacaagctgaaaaagctgatcaacaagagcc
ccgaaaagctgctgatgtaccaccacgacccccagacctaccagaaactgaagctgattatggaacagtacggcgacgagaagaatccc
ctgtacaagtactacgaggaaaccgggaactacctgaccaagtactccaaaaaggacaacggccccgtgatcaagaagattaagtattac
ggcaacaaactgaacgcccatctggacatcaccgacgactaccccaacagcagaaacaaggtcgtgaagctgtccctgaagccctacag
attcgacgtgtacctggacaatggcgtgtacaagttcgtgaccgtgaagaatctggatgtgatcaaaaaagaaaactactacgaagtgaata
gcaagtgctatgaggaagctaagaagctgaagaagatcagcaaccaggccgagtttatcgcctccttctacaacaacgatctgatcaagat
caacggcgagctgtatagagtgatcggcgtgaacaacgacctgctgaaccggatcgaagtgaacatgatcgacatcacctaccgcgagta
cctggaaaacatgaacgacaagaggcccccccaggatcattaagacaatcgcctccaagacccagagcattaagaagtacagcacagaca
ttctgggcaacctgtatgaagtgaaatctaagaagcaccctcagatcatcaaaaagggcaaaaggccggcggccacgaaaaaggcGcT
CGAGGGCGGAGGCGGGAGCGGATCCCCCTCCCGGCTCCAGATGttcttcgctaataaccacgacc
aggaatttgaccctccaaaggtttacccacctgtcccagctgagaagaggaagcccatccgggtgctgtctctctttgatggaatcgctacag
ggctcctggtgctgaaggacttgggcattcaggtggaccgctacattgcctcggaggtgtgtgaggactccatcacggtgggcatggtgcg

gcaccaggggaagatcatgtacgtcggggacgtccgcagcgtcacacagaagcatatccaggagtggggcccattcgatctggtgattg
ggggcagtccctgcaatgacctctccatcgtcaaccctgctcgcaagggcctctacgagggcactggccggctcttctttgagttctaccgc
ctcctgcatgatgcgcggcccaaggagggagatgatcgccccttcttctggctctttgagaatgtggtggccatgggcgttagtgacaagag
ggacatctcgcgatttctcgagtccaaccctgtgatgattgatgccaaagaagtgtcagctgcacacagggcccgctacttctggggtaacct
tcccggtatgaacaggccgttggcatccactgtgaatgataagctggagctgcaggagtgtctggagcatggcaggatagccaagttcagc
aaagtgaggaccattactacgaggtcaaactccataaagcagggcaaaGACCAGCATTTTCCTGTGTTCATGAAT
GAGAAAGAGgacatcttatggtgcactgaaatggaaagggtatttggtttcccagtccactatactgacgtgtccaacatgagccgct
tggcgaggcagagactgctgggccggtcatggagcgtgccagtcatccgccacctcttcgctcCGCTGAAGGAGTATTTT
GCGTGTGTGtccggccggcccggatccGGCGCAACAAACTTCTCTCTGCTGAAACAAGCCGGA
GATGTCGAAGAGAATCCTGGACCGACCGAGTACAAGCCCACGGTGCGCCTCGCCAC
CCGCGACGACGTCCCCAGGGCCGTACGCACCCTCGCCGCCGCGTTCGCCGACTACCC
CGCCACGCGCCACACCGTCGATCCGGACCGCCACATCGAGCGGGTCACCGAGCTGC
AAGAACTCTTCCTCACGCGCGTCGGGCTCGACATCGGCAAGGTGTGGGTCGCGGAC
GACGGCGCCGCGGTGGCGGTCTGGACCACGCCGGAGAGCGTCGAAGCGGGGGCGG
TGTTCGCCGAGATCGGCCCGCGCATGGCCGAGTTGAGCGGTTCCCGGCTGGCCGCGC
AGCAACAGATGGAAGGCCTCCTGGCGCCGCACCGGCCCAAGGAGCCCGCGTGGTTC
CTGGCCACCGTCGGAGTCTCGCCCGACCACCAGGGCAAGGGTCTGGGCAGCGCCGT
CGTGCTCCCCGGAGTGGAGGCGGCCGAGCGCGCCGGGGTGCCCGCCTTCCTGGAGA
CCTCCGCGCCCCGCAACCTCCCCTTCTACGAGCGGCTCGGCTTCACCGTCACCGCCG
ACGTCGAGGTGCCCGAAGGACCGCGCACCTGGTGCATGACCCGCAAGCCCGGTGCC
TGAACGCGTTAAGTCGACAATCAACCTCTGGATTACAAAATTTGTGAAAGATTGACT
GGTATTCTTAACTATGTTGCTCCTTTTACGCTATGTGGATACGCTGCTTTAATGCCTT
TGTATCATGCTATTGCTTCCCGTATGGCTTTCATTTTCTCCTCCTTGTATAAATCCTGG
TTGCTGTCTCTTTATGAGGAGTTGTGGCCCGTTGTCAGGCAACGTGGCGTGGTGTGC
ACTGTGTTTGCTGACGCAACCCCCACTGGTTGGGGCATTGCCACCACCTGTCAGCTC
CTTTCCGGGACTTTCGCTTTCCCCCTCCCTATTGCCACGGCGGAACTCATCGCCGCCT
GCCTTGCCCGCTGCTGGACAGGGGCTCGGCTGTTGGGCACTGACAATTCCGTGGTGT
TGTCGGGGAAATCATCGTCCTTTCCTTGGCTGCTCGCCTGTGTTGCCACCTGGATTCT
GCGCGGGACGTCCTTCTGCTACGTCCCTTCGGCCCTCAATCCAGCGGACCTTCCTTCC
CGCGGCCTGCTGCCGGCTCTGCGGCCTCTTCCGCGTCTTCGCCTTCGCCCTCAGACG
AGTCGGATCTCCCTTTGGGCCGCCTCCCCGCGTCGACTTTAAGACCAATGACTTACA
AGGCAGCTGTAGATCTTAGCCACTTTTTAAAAGAAAAGGGGGGACTGGAAGGGCTA
ATTCACTCCCAACGAAGACAAGATCTGCTTTTTGCTTGTACTGGGTCTCTCTGGTTAG
ACCAGATCTGAGCCTGGGAGCTCTCTGGCTAACTAGGGAACCCACTGCTTAAGCCTC
AATAAAGCTTGCCTTGAGTGCTTCAAGTAGTGTGTGCCCGTCTGTTGTGTGACTCTG
GTAACTAGAGATCCCTCAGACCCTTTTAGTCAGTGTGGAAAATCTCTAGCAGggcccgttt
aaacccgctgatcagcctcgactgtgccttctagttgccagccatctgttgtttgcccctcccccgtgccttccttgaccctggaaggtgccact
cccactgtcctttcctaataaaatgaggaaattgcatcgcattgtctgagtaggtgtcattctattctggggggtggggtggggcaggacagc
aagggggaggattgggaagacaatagcaggcatgctggggatgcggtgggctctatggcttctgaggcggaaagaaccagctggggct
ctagggggtatccccacgcgccctgtagcggcgcattaagcgcggcgggtgtggtggttacgcgcagcgtgaccgctacacttgccagc
gccctagcgcccgctcctttcgctttcttcccttcctttctcgccacgttcgccggctttccccgtcaagctctaaatcgggggctccctttaggg
ttccgatttagtgctttacggcacctcgaccccaaaaaacttgattagggtgatggttcacgtagtgggccatcgccctgatagacggtttttcg
ccctttgacgttggagtccacgttctttaatagtggactcttgttccaaactggaacaacactcaaccctatctcggtctattcttttgatttataatg
gttacaaataaagcaatagcatcacaaatttcacaaataaagcatttttttcactgcattctagttgtggtttgtccaaactcatcaatgtatcttatc
atgtctgtataccgtcgacctctagctagagcttggcgtaatcatggtcatagctgtttcctgtgtgaaattgttatccgctcacaattccacacaa
catacgagccggaagcataaagtgtaaagcctggggtgcctaatgagtgagctaactcacattaattgcgttgcgctcactgcccgctttcca

gtcgggaaacctgtcgtgccagctgcattaatgaatcggccaacgcgcggggagaggcggtttgcgtattgggcgctcttccgcttcctcg
ctcactgactcgctgcgctcggtcgttcggctgcggcgagcggtatcagctcactcaaaggcggtaatacggttatccacagaatcagggg
ataacgcaggaaagaacatgtgagcaaaaggccagcaaaaggccaggaaccgtaaaaaggccgcgttgctggcgtttttccataggctc
cgcccccctgacgagcatcacaaaaatcgacgctcaagtcagaggtggcgaaacccgacaggactataaagataccaggcgtttccccct
ggaagctccctcgtgcgctctcctgttccgaccctgccgcttaccggatacctgtccgcctttctcccttcgggaagcgtggcgctttctcata
gctcacgctgtaggtatctcagttcggtgtaggtcgttcgctccaagctgggctgtgtgcacgaaccccccgttcagcccgaccgctgcgcc
ttatccggtaactatcgtcttgagtccaacccggtaagacacgacttatcgccactggcagcagccactggtaacaggattagcagagcgag
gtatgtaggcggtgctacagagttcttgaagtggtggcctaactacggctacactagaagaacagtatttggtatctgcgctctgctgaagcc
agttaccttcggaaaaagagttggtagctcttgatccggcaaacaaaccaccgctggtagcggtggtttttttgtttgcaagcagcagattacg
cgcagaaaaaaaggatctcaagaagatcctttgatcttttctacggggtctgacgctcagtggaacgaaaactcacgttaagggattttggtc
atgagattatcaaaaaggatcttcacctagatccttttaaattaaaaatgaagttttaaatcaatctaaagtatatatgagtaaacttggtctgacag
ttaccaatgcttaatcagtgaggcacctatctcagcgatctgtctatttcgttcatccatagttgcctgactccccgtcgtgtagataactacgata
cgggagggcttaccatctggccccagtgctgcaatgataccgcgagacccacgctcaccggctccagatttatcagcaataaaccagcca
gccggaagggccgagcgcagaagtggtcctgcaactttatccgcctccatccagtctattaattgttgccgggaagctagagtaagtagttc
gccagttaatagtttgcgcaacgttgttgccattgctacaggcatcgtggtgtcacgctcgtcgtttggtatggcttcattcagctccggttccca
acgatcaaggcgagttacatgatcccccatgttgtgcaaaaaagcggttagctccttcggtcctccgatcgttgtcagaagtaagttggccgc
agtgttatcactcatggttatggcagcactgcataattctcttactgtcatgccatccgtaagatgcttttctgtgactggtgagtactcaaccaag
tcattctgagaatagtgtatgcggcgaccgagttgctcttgcccggcgtcaatacgggataataccgcgccacatagcagaactttaaaagtg
ctcatcattggaaaacgttcttcggggcgaaaactctcaaggatcttaccgctgttgagatccagttcgatgtaacccactcgtgcacccaact
gatcttcagcatcttttactttcaccagcgtttctgggtgagcaaaaacaggaaggcaaaatgccgcaaaaaagggaataagggcgacacg
gaaatgttgaatactcatactcttcctttttcaatattattgaagcatttatcagggttattgtctcatgagcggatacatatttgaatgtatttagaaa
aataaacaaataggggttccgcgcacatttccccgaaaagtgccacctgacgtcgacggatcgggagatctcccgatcccctatggtgcac
tctcagtacaatctgctctgatgccgcatagttaagccagtatctgctccctgcttgtgtgttggaggtcgctgagtagtgcgcgagcaaaattt
aagctacaacaaggcaaggcttgaccgacaattgcatgaagaatctgcttagggttaggcgttttgcgctgcttcgcgatgtacgggccaga
tatacgcgctgacattgattatt

### 7. BK1837- pLV-CMV-SNCAintron1-AlphaSyn-d2EGFP-P2A-NLuc-SV40-Puro-WPRE (pCL35)

gcacttttcggggaaatgtgcgcggaacccctatttgtttatttttctaaatacattcaaatatgtatccgctcatgagacaataaccctgataaat
gcttcaataatattgaaaaaggaagagtatgagtattcaacatttccgtgtcgcccttattcccttttttgcggcattttgccttcctgtttttgctcac
ccagaaacgctggtgaaagtaaaagatgctgaagatcagttgggtgcacgagtgggttacatcgaactggatctcaacagcggtaagatcc
ttgagagttttcgccccgaagaacgttttccaatgatgagcacttttaaagttctgctatgtggcgcggtattatcccgtattgacgccgggcaa
gagcaactcggtcgccgcatacactattctcagaatgacttggttgagtactcaccagtcacagaaaagcatcttacggatggcatgacagta
agagaattatgcagtgctgccataaccatgagtgataacactgcggccaacttacttctgacaacgatcggaggaccgaaggagctaaccg
cttttttgcacaacatgggggatcatgtaactcgccttgatcgttgggaaccggagctgaatgaagccataccaaacgacgagcgtgacacc
acgatgcctgtagcaatggcaacaacgttgcgcaaactattaactggcgaactacttactctagcttcccggcaacaattaatagactggatg
gaggcggataaagttgcaggaccacttctgcgctcggcccttccggctggctggtttattgctgataaatctggagccggtgagcgtgggtc
tcgcggtatcattgcagcactggggccagatggtaagccctcccgtatcgtagttatctacacgacggggagtcaggcaactatggatgaa
cgaaatagacagatcgctgagataggtgcctcactgattaagcattggtaactgtcagaccaagtttactcatatatactttagattgatttaaaa
cttcatttttaatttaaaaggatctaggtgaagatcctttttgataatctcatgaccaaaatcccttaacgtgagttttcgttccactgagcgtcaga
ccccgtagaaaagatcaaaggatcttcttgagatcctttttttctgcgcgtaatctgctgcttgcaaacaaaaaaaccaccgctaccagcggtg
gtttgtttgccggatcaagagctaccaactctttttccgaaggtaactggcttcagcagagcgcagataccaaatactgttcttctagtgtagcc
gtagttaggccaccacttcaagaactctgtagcaccgcctacatacctcgctctgctaatcctgttaccagtggctgctgccagtggcgataa

gtcgtgtcttaccgggttggactcaagacgatagttaccggataaggcgcagcggtcgggctgaacggggggttcgtgcacacagcccag
cttggagcgaacgacctacaccgaactgagatacctacagcgtgagctatgagaaagcgccacgcttcccgaagggagaaaggcggac
aggtatccggtaagcggcagggtcggaacaggagagcgcacgagggagcttccagggggaaacgcctggtatctttatagtcctgtcgg
gtttcgccacctctgacttgagcgtcgatttttgtgatgctcgtcaggggggcggagcctatggaaaaacgccagcaacgcggcctttttacg
gttcctggccttttgctggccttttgctcacatgttctttcctgcgttatcccctgattctgtggataaccgtattaccgcctttgagtgagctgatac
cgctcgccgcagccgaacgaccgagcgcagcgagtcagtgagcgaggaagcggaagagcgcccaatacgcaaaccgcctctccccg
cgcgttggccgattcattaatgcagctggcacgacaggtttcccgactggaaagcgggcagtgagcgcaacgcaattaatgtgagttagct
cactcattaggcaccccaggctttacactttatgcttccggctcgtatgttgtgtggaattgtgagcggataacaatttcacacaggaaacagct
atgaccatgattacgccaagcgcgcaattaaccctcactaaagggaacaaaagctggagctgcaagcttaatgtagtcttatgcaatactctt
gtagtcttgcaacatggtaacgatgagttagcaacatgccttacaaggagagaaaaagcaccgtgcatgccgattggtggaagtaaggtgg
tacgatcgtgccttattaggaaggcaacagacgggtctgacatggattggacgaaccactgaattgccgcattgcagagatattgtatttaagt
gcctagctcgatacataaacgggtctctctggttagaccagatctgagcctgggagctctctggctaactagggaacccactgcttaagcctc
aataaagcttgccttgagtgcttcaagtagtgtgtgcccgtctgttgtgtgactctggtaactagagatccctcagacccttttagtcagtgtgga
aaatctctagcagtggcgcccgaacagggacttgaaagcgaaagggaaaccagaggagctctctcgacgcaggactcggcttgctgaag
cgcgcacggcaagaggcgaggggcggcgactggtgagtacgccaaaaattttgactagcggaggctagaaggagagagatgggtgcg
agagcgtcagtattaagcgggggagaattagatcgcgatgggaaaaaattcggttaaggccagggggaaagaaaaaatataaattaaaac
atatagtatgggcaagcagggagctagaacgattcgcagttaatcctggcctgttagaaacatcagaaggctgtagacaaatactgggaca
gctacaaccatcccttcagacaggatcagaagaacttagatcattatataatacagtagcaaccctctattgtgtgcatcaaaggatagagata
aaagacaccaaggaagctttagacaagatagaggaagagcaaaacaaaagtaagaccaccgcacagcaagcggccgctgatcttcaga
cctggaggaggagatatgagggacaattggagaagtgaattatataaatataaagtagtaaaaattgaaccattaggagtagcacccaccaa
ggcaaagagaagagtggtgcagagagaaaaaagagcagtgggaataggagctttgttccttgggttcttgggagcagcaggaagcactat
gggcgcagcgtcaatgacgctgacggtacaggccagacaattattgtctggtatagtgcagcagcagaacaatttgctgagggctattgag
gcgcaacagcatctgttgcaactcacagtctggggcatcaagcagctccaggcaagaatcctggctgtggaaagatacctaaaggatcaa
cagctcctggggatttggggttgctctggaaaactcatttgcaccactgctgtgccttggaatgctagttggagtaataaatctctggaacagat
ttggaatcacacgacctggatggagtgggacagagaaattaacaattacacaagcttaatacactccttaattgaagaatcgcaaaaccagc
aagaaaagaatgaacaagaattattggaattagataaatgggcaagtttgtggaattggtttaacataacaaattggctgtggtatataaaattat
tcataatgatagtaggaggcttggtaggtttaagaatagtttttgctgtactttctatagtgaatagagttaggcagggatattcaccattatcgttt
cagacccacctcccaaccccgaggggacccgacaggcccgaaggaatagaagaagaaggtggagagagagacagagacagatccat
tcgattagtgaacggatctcgacggtatcggttaacttttaaaagaaaaggggggattggggggtacagtgcaggggaaagaatagtagac
ataatagcaacagacatacaaactaaagaattacaaaaacaaattacaaaattcaaaattttatCGATTTAATTAACGCGTTA
CATAACTTACGGTAAATGGCCCGCCTGGCTGACCGCCCAACGACCCCCGCCCATTGAC
GTCAATAATGACGTATGTTCCCATAGTAACGCCAATAGGGACTTTCCATTGACGTCAAT
GGGTGGAGTATTTACGGTAAACTGCCCACTTGGCAGTACATCAAGTGTATCATATGCC
AAGTACGCCCCCTATTGACGTCAATGACGGTAAATGGCCCGCCTGGCATTATGCCCAG
TACATGACCTTATGGGACTTTCCTACTTGGCAGTACATCTACGTATTAGTCATCGCTATT
ACCATGGTCGAGGTGAGCCCCACGTTCTGCTTCACTCTCCCCATCTCCCCCCCCTCCC
CACCCCCAATTTTGTATTTATTTATTTTTTAATTATTTTGTGCAGCGATGGGGGCGGGGG
GGGGGGGGGGCGCGCGCCAGGCGGGGCGGGGCGGGGCGAGGGGCGGGGCGGGG
CGAGGCGGAGAGGTGCGGCGGCAGCCAATCAGAGCGGCGCGCTCCGAAAGTTTCCT
TTTATGGCGAGGCGGCGGCGGCGGCGGCCCTATAAAAAGCGAAGCGCGCGGCGGGC

Gcgggcgtctttggaaatcctggagaacgccggatgggagacgaatggtcgtgggcaccgggagggggtggtgctgccatgaggacc
cgctgggccaggtctctgggaggtgagtacttgtccctttggggagcctaaggaaagagacttgacctggctttcgtcctgcttctgatattcc
cttctccacaagggctgagagattaggctgcttctccgggatccgcttttccccgggaaacgcgaggatgctccatggagcgtgagcatcc
aacttttctctcacataaaatctgtctgcccgctctcttggtttttctctgtaaagtaagcaagctgcgtttggcaaataatgaaatggaagtgcaa
ggaggccaagtcaacaggtggtaacgggttaacaagtgctggcgcggggtccgctagggtggaggctgagaacgccccctcgggtggc
tggcgcggggttggagacggcccgcgagtgtgagcggcgcctgctcagggtagatagctgagggcggATGgatgtattcatgaaagg
actttcaaaggccaaggagggagttgtggctgctgctgagaaaaccaaacagggtgtggcagaagcagcaggaaagacaaaagagggt
gttctctatgtaggctccaaaaccaaggagggagtggtgcatggtgtggcaacagtggctgagaagaccaaagagcaagtgacaaatgtt
ggaggagcagtggtgacgggtgtgacagcagtagcccagaagacagtggagggagcagggagcattgcagcagccactggctttgtca
aaaaggaccagttgggcaagaatgaagaaggagccccacaggaaggaattctggaagatatgcctgtggatcctgacaatgaggcttatg
aaatgccttctgaggaagggtatcaagactacgaacctgaagAGGAGGAGAGTGACTTAATTAAAccggtcgccacc
atggtgagcaagggcgaggagctgttcaccggggtggtgcccatcctggtcgagctggacggcgacgtaaacggccacaagttcagcgt
gtccggcgagggcgagggcgatgccacctacggcaagctgaccctgaagttcatctgcaccaccggcaagctgcccgtgccctggccc
accctcgtgaccaccctgacctacggcgtgcagtgcttcagccgctaccccgaccacatgaagcagcacgacttcttcaagtccgccatgc
ccgaaggctacgtccaggagcgcaccatcttcttcaaggacgacggcaactacaagacccgcgccgaggtgaagttcgagggcgacac
cctggtgaaccgcatcgagctgaagggcatcgacttcaaggaggacggcaacatcctggggcacaagctggagtacaactacaacagcc
acaacgtctatatcatggccgacaagcagaagaacggcatcaaggtgaacttcaagatccgccacaacatcgaggacggcagcgtgcag
ctcgccgaccactaccagcagaacacccccatcggcgacggccccgtgctgctgcccgacaaccactacctgagcacccagtccgccct
gagcaaagaccccaacgagaagcgcgatcacatggtcctgctggagttcgtgaccgccgccgggatcactctcggcatggacgagctgt
acaagaagcttagccatggcttcccgccggaggtggaggagcaggatgatggcacgctgcccatgtcttgtgcccaggagagcgggatg
gaccgtcaccctgcagcctgtgcttctgctaggatcaatgtgGaggAggGcgGCTCgAGAGCCACGAACTTCTCT
CTGTTAAAGCAAGCAGGAGACGTGGAAGAAAACCCCGGTCCCGCCACCATGGTCTT
CACACTCGAAGATTTCGTTGGGGACTGGCGACAGACAGCCGGCTACAACCTGGACC
AAGTCCTTGAACAGGGAGGTGTGTCCAGTTTGTTTCAGAATCTCGGGGTGTCCGTAA
CTCCGATCCAAAGGATTGTCCTGAGCGGTGAAAATGGGCTGAAGATCGACATCCATG
TCATCATCCCGTATGAAGGTCTGAGCGGCGACCAAATGGGCCAGATCGAAAAAATTTT
TAAGGTGGTGTACCCTGTGGATGATCATCACTTTAAGGTGATCCTGCACTATGGCACA
CTGGTAATCGACGGGGTTACGCCGAACATGATCGACTATTTCGGACGGCCGTATGAAG
GCATCGCCGTGTTCGACGGCAAAAAGATCACTGTAACAGGGACCCTGTGGAACGGC
AACAAAATTATCGACGAGCGCCTGATCAACCCCGACGGCTCCCTGCTGTTCCGAGTA
ACCATCAACGGAGTGACCGGCTGGCGGCTGTGCGAACGCATTCTGGCGTAAGtcgaCcat
gcatctagacccagcggCgCGcCCCTGTGGAATGTGTGTCAGTTAGGGTGTGGAAAGTCCCCAG
GCTCCCCAGCAGGCAGAAGTATGCAAAGCATGCATCTCAATTAGTCAGCAACCAGGT
GTGGAAAGTCCCCAGGCTCCCCAGCAGGCAGAAGTATGCAAAGCATGCATCTCAATT
AGTCAGCAACCATAGTCCCGCCCCTAACTCCGCCCATCCCGCCCCTAACTCCGCCCAG
TTCCGCCCATTCTCCGCCCCATGGCTGACTAATTTTTTTTATTTATGCAGAGGCCGAGG
CCGCCTCGGCCTCTGAGCTATTCCAGAAGTAGTGAGGAGGCTTTTTTGGAGGCCTAG
GCTTTTGCAAAAAGCTTACCATGACCGAGTACAAGCCCACGGTGCGCCTCGCCACCC
GCGACGACGTCCCCAGGGCCGTACGCACCCTCGCCGCCGCGTTCGCCGACTACCCCG
CCACGCGCCACACCGTCGATCCGGACCGCCACATCGAGCGGGTCACCGAGCTGCAA

GAACTCTTCCTCACGCGCGTCGGGCTCGACATCGGCAAGGTGTGGGTCGCGGACGAC
GGCGCCGCGGTGGCGGTCTGGACCACGCCGGAGAGCGTCGAAGCGGGGGCGGTGTT
CGCCGAGATCGGCCCGCGCATGGCCGAGTTGAGCGGTTCCCGGCTGGCCGCGCAGC
AACAGATGGAAGGCCTCCTGGCGCCGCACCGGCCCAAGGAGCCCGCGTGGTTCCTG
GCCACCGTCGGCGTCTCGCCCGACCACCAGGGCAAGGGTCTGGGCAGCGCCGTCGT
GCTCCCCGGAGTGGAGGCGGCCGAGCGCGCCGGGGTGCCCGCCTTCCTGGAGACCT
CCGCGCCCCGCAACCTCCCCTTCTACGAGCGGCTCGGCTTCACCGTCACCGCCGACG
TCGAGGTGCCCGAAGGACCGCGCACCTGGTGCATGACCCGCAAGCCCGGTGCCTGAc
gGgcgcgccgAATTCGACAATCAACCTCTGGATTACAAAATTTGTGAAAGATTGACTGGT
ATTCTTAACTATGTTGCTCCTTTTACGCTATGTGGATACGCTGCTTTAATGCCTTTGTAT
CATGCTATTGCTTCCCGTATGGCTTTCATTTTCTCCTCCTTGTATAAATCCTGGTTGCTG
TCTCTTTATGAGGAGTTGTGGCCCGTTGTCAGGCAACGTGGCGTGGTGTGCACTGTGT
TTGCTGACGCAACCCCCACTGGTTGGGGCATTGCCACCACCTGTCAGCTCCTTTCCG
GGACTTTCGCTTTCCCCCTCCCTATTGCCACGGCGGAACTCATCGCCGCCTGCCTTGC
CCGCTGCTGGACAGGGGCTCGGCTGTTGGGCACTGACAATTCCGTGGTGTTGTCGGG
GAAATCATCGTCCTTTCCTTGGCTGCTCGCCTGTGTTGCCACCTGGATTCTGCGCGGG
ACGTCCTTCTGCTACGTCCCTTCGGCCCTCAATCCAGCGGACCTTCCTTCCCGCGGCC
TGCTGCCGGCTCTGCGGCCTCTTCCGCGTCTTCGCCTTCGCCCTCAGACGAGTCGGAT
CTCCCTTTGGGCCGCCTCCCCGCGTCGACTTTAAGACCAATGACTTACAAGGTACctttaa
gaccaatgacttacaaggcagctgtagatcttagccactttttaaaagaaaaggggggactggaagggctaattcactcccaacgaagaca
agatctgctttttgcttgtactgggtctctctggttagaccagatctgagcctgggagctctctggctaactagggaacccactgcttaagcctc
aataaagcttgccttgagtgcttcaagtagtgtgtgcccgtctgttgtgtgactctggtaactagagatccctcagacccttttagtcagtgtgga
aaatctctagcagtagtagttcatgtcatcttattattcagtatttataacttgcaaagaaatgaatatcagagagtgagaggaacttgtttattgca
gcttataatggttacaaataaagcaatagcatcacaaatttcacaaataaagcatttttttcactgcattctagttgtggtttgtccaaactcatcaat
gtatcttatcatgtctggctctagctatcccgcccctaactccgcccatcccgcccctaactccgcccagttccgcccattctccgccccatgg
ctgactaattttttttatttatgcagaggccgaggccgcctcggcctctgagctattccagaagtagtgaggaggcttttttggaggcctaggga
cgtacccaattcgccctatagtgagtcgtattacgcgcgctcactggccgtcgttttacaacgtcgtgactgggaaaaccctggcgttaccca
acttaatcgccttgcagcacatccccctttcgccagctggcgtaatagcgaagaggcccgcaccgatcgcccttcccaacagttgcgcagc
ctgaatggcgaatgggacgcgccctgtagcggcgcattaagcgcggcgggtgtggtggttacgcgcagcgtgaccgctacacttgccag
cgccctagcgcccgctcctttcgctttcttcccttcctttctcgccacgttcgccggctttccccgtcaagctctaaatcgggggctccctttagg
gttccgatttagtgctttacggcacctcgaccccaaaaaacttgattagggtgatggttcacgtagtgggccatcgccctgatagacggtttttc
gccctttgacgttggagtccacgttctttaatagtggactcttgttccaaactggaacaacactcaaccctatctcggtctattcttttgatttataa
gggattttgccgatttcggcctattggttaaaaaatgagctgatttaacaaaaatttaacgcgaattttaacaaaatattaacgcttacaatttaggt
g
